# Predicting potential impacts of ocean acidification on marine calcifiers from the Southern Ocean

**DOI:** 10.1101/2020.11.15.384131

**Authors:** Blanca Figuerola, Alyce M. Hancock, Narissa Bax, Vonda Cummings, Rachel Downey, Huw J. Griffiths, Jodie Smith, Jonathan S. Stark

**Author notes:** Correspondence: Dr. Blanca Figuerola.

## Abstract

Understanding the vulnerability of marine calcifiers to ocean acidification is a critical issue, especially in the Southern Ocean (SO), which is likely to be the one of the first, and most severely affected regions. Since the industrial revolution, ~30% of anthropogenic CO_2_ has been absorbed by the oceans. Seawater pH levels have already decreased by 0.1 and are predicted to decline by ~ 0.3 by the year 2100. This process, known as ocean acidification (OA), is shallowing the saturation horizon, which is the depth below which calcium carbonate (CaCO_3_) dissolves, likely increasing the vulnerability of many marine calcifiers to dissolution. The negative impact of OA may be seen first in species depositing more soluble CaCO_3_ mineral phases such as aragonite and high-Mg calcite (HMC). These negative effects may become even exacerbated by increasing sea temperatures. Here we combine a review and a quantitative meta-analysis to provide an overview of the current state of knowledge about skeletal mineralogy of major taxonomic groups of SO marine calcifiers and to make predictions about how OA might affect different taxa. We consider their geographic range, skeletal mineralogy, biological traits and potential strategies to overcome OA. The meta-analysis of studies investigating the effects of the OA on a range of biological responses such as shell state, development and growth rate shows response variation depending on mineralogical composition. Species-specific responses due to mineralogical composition suggest taxa with calcitic, aragonitic and HMC skeletons may be more vulnerable to the expected carbonate chemistry alterations, and low magnesium calcite (LMC) species may be mostly resilient. Environmental and biological control on the calcification process and/or Mg content in calcite, biological traits and physiological processes are also expected to influence species specific responses.

## 1 Introduction

Since the industrial revolution, the concentration of carbon dioxide (CO_2_) released to the atmosphere has increased from 280 to above 400 μatm due to human activities such as burning of fossil fuels and deforestation. Approximately 30% of this has been absorbed by the oceans (Feely et al., 2004). Seawater pH levels have already decreased by 0.1 and are predicted to decline by ~ 0.3 (851-1370 μatm) by the year 2100 (IPCC, 2014, 2019). This process, known as ocean acidification (OA), also lowers both carbonate ion concentration ([CO_3_^2−^]) and saturation state (Ω) of seawater with respect to calcium carbonate (CaCO_3_) minerals. In particular, Ω is a function of [CO_3_^2−^] and calcium ion concentrations ([Ca^2+^]) expressed as

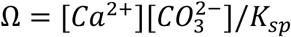

where K_sp_ is the solubility product, which depends on the specific CaCO_3_ mineral phase, temperature, salinity, and pressure (Zeebe, 2012).

Surface waters are naturally supersaturated with respect to carbonate (Ω > 1), where the highest [CO_3_^2−^] is found as a result of surface photosynthesis. The [CO_3_^2−^] decreases, and the solubility of CaCO_3_ minerals (such as aragonite) increases, with depth due to the higher pressure and lower temperature. Thus, Ω is higher in shallow, warm waters than in cold waters and at depth (Feely et al., 2004). Current aragonite saturation in Southern Ocean (SO) surface waters ranges from Ω^arag^ of 1.22 to 2.42, compared with values of 3.5 to 4 in the tropics (Jiang et al., 2015), and varies spatially, e.g. across ocean basins, and seasonally. However, OA is both decreasing saturation levels and shallowing the carbonate saturation horizon, which is the depth below which CaCO_3_ can dissolve (Ω < 1). This is likely to increase the vulnerability of CaCO_3_ sediments and many marine calcifiers (CaCO_3_ shell/skeleton-building organisms) to dissolution (Fig. 1) (Haese et al., 2014). As CaCO_3_ sediments have important roles in global biogeochemical cycling and provide suitable habitat for many marine calcifying species, undersaturated seawater conditions will also lead to unprecedented challenges and alterations to the function, structure and distribution of carbonate ecosystems exposed to these conditions (Andersson et al., 2008).

**Fig. 1.**
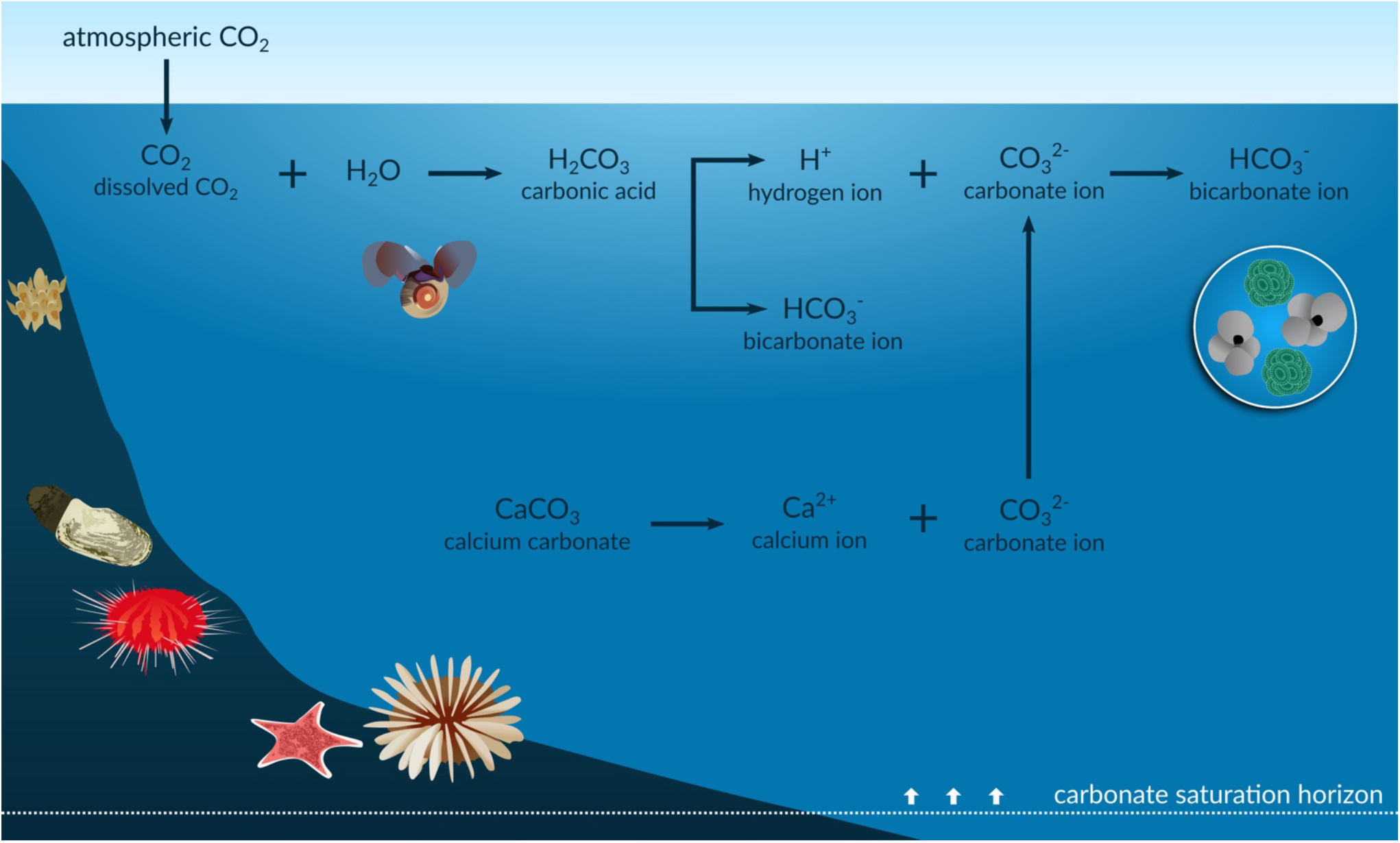
Infographic of the ocean acidification process. The anthropogenic CO_2_ absorbed by the oceans results in an increase in the concentration of hydrogen ions (H^+^) and in bicarbonate ions (HCO_3_^−^) and a decrease in carbonate ions (CO_3_^2−^). The reduction in CO_3_^2−^ is shallowing the carbonate saturation horizons, with potential impacts on shells and skeletons of marine calcifiers such as foraminifera, corals, echinoderms, molluscs and bryozoans.

A number of marine calcifiers, particularly benthic populations from the tropics and deep waters, are already exposed to undersaturated conditions with respect to their species-specific mineralogies (Lebrato et al., 2016). This suggests some organisms have evolved compensatory physiological mechanisms to maintain the calcification process (Lebrato et al., 2016). This does not mean that these taxa will cope with sudden decreases in the Ω, rather it illustrates the complexity of taxon-level responses. For instance, SO marine calcifiers may be particularly sensitive to these changes since the SO is characterized by much lower natural variability in surface ocean [CO_3_^2−^] (Conrad and Lovenduski, 2015). The processes responsible for making calcifying organisms vulnerable to OA have been much discussed. Recent research suggests that the reduction in seawater pH mainly drives changes in calcification rates (the production and deposition of CaCO_3_), rather than the reduction in [CO_3_^2−^] (Cyronak et al., 2016), with a predicted drop in calcification rates by 30 to 40% by mid-century due to OA (Kleypas et al., 1999). The long-term persistence of calcifying taxa depends on calcification exceeding the loss of CaCO_3_ that occurs through breakdown, export and dissolution processes (Andersson and Gledhill, 2013). Therefore, even though species-specific responses to OA are expected, a tendency toward reduction in overall CaCO_3_ production at the local or community level is anticipated (Andersson and Gledhill, 2013) with potentially adverse implications across SO ecosystems.

### 1.1 Southern Ocean acidification

The Southern Ocean, which covers about 34.8 million km^2^, is widely anticipated to be the one of the first, and most severely affected region from OA due to naturally low levels of CaCO_3_, the increased solubility of CO_2_ at low temperatures, and a lower buffering capacity (Orr et al., 2005). Thus, understanding the vulnerability of Antarctic marine calcifiers to OA is a critical issue, particularly in Antarctic Peninsula, which is also one of the fastest warming areas on Earth (Vaughan and Marshall, 2003). To establish the potential threat of OA to SO marine calcifiers, it is crucial to consider different factors that may influence the vulnerability of their skeletons to OA. These include interacting factors (e.g. warming), the local seasonal and spatial variability in the carbonate system, the distribution of sedimentary CaCO_3_ (which provides suitable habitat for many benthic calcifiers), and a range of different phyla and species with diverse geographic range, skeletal mineralogies, biological traits, physiological processes and strategies to combat pH changes.

#### 1.1.1 Local seasonal and spatial variability in carbonate system

The depth of the present-day SO Aragonite Saturation Horizon (ASH) exceeds 1,000 m across most of the basin, while naturally shallower saturation horizons (~400 m) occur in the core of the Antarctic Circumpolar Current due to upwelling of CO_2_-rich deep water (Negrete-García et al., 2019). The Calcite Saturation Horizon (CSH) is much deeper (> 2000 m in some parts of the SO) than the ASH, since aragonite is more soluble than calcite (Feely et al., 2004; Barnes and Peck, 2008). It is thus predicted that the ASH will reach surface waters earlier than the CSH (McNeil and Matear, 2008). In particular, it is anticipated that aragonite in some SO regions (south of 60°S) will be undersaturated by 2050 (Orr et al., 2005; McNeil and Matear, 2008) and that 70% of the SO could be undersaturated by 2100 (Hauri et al., 2016). OA will drive even the least soluble form of calcium carbonate, calcite, to undersaturation by 2095 in some SO waters (McNeil and Matear, 2008). However, the depth and year of emergence of a shallow ASH can vary spatially due to natural variation in the present-day saturation horizon depth and in the physical circulation of the SO (Negrete-García et al., 2019).

Aragonite undersaturation in surface waters is predicted to occur initially during the austral winter (Conrad and Lovenduski, 2015; Negrete-García et al., 2019) as the [CO_3_^2−^] of surface waters decreases south of the Polar Front due to the strong persistent winter winds that promote the upwelling of carbonate-depleted deep waters (Orr et al., 2005). In addition, lower temperatures enhance uptake of atmospheric CO_2_ (Orr et al., 2005). Specifically, several authors (McNeil and Matear, 2008; McNeil et al., 2010; Mattsdotter Björk et al., 2014) predicted that wintertime aragonite undersaturation will occur by 2030 in the latitudinal band between 65 and 70°S where deep-water upwelling occurs. Recently, Negrete-García et al. (2019) considered that the rate of change, and therefore the onset of aragonite undersaturation, was occurring more rapidly than previously projected. These predicted changes imply there will be a sudden decline in the availability of suitable habitat for diverse SO taxa such as pteropods, foraminifera, cold-water corals and coralline algae (Negrete-García et al., 2019).

#### 1.1.2 Distribution of carbonate sediments

The distribution of sedimentary CaCO_3_ on the Antarctic shelves displays distinct regional and depth patterns (Hauck et al., 2012). This is evident from the deposition and preservation of carbonates in the surface sediments. These are driven by the flux of organic matter to the ocean floor (related to primary production) and its respiration and remineralization in the sediments, transport of carbonate material by currents, and Ω of the water mass above the sediment (Hauck et al., 2012). In particular, a higher flux of organic matter to the seafloor increases respiration in sediments and alters carbonate chemistry. CaCO_3_ contents of shelf surface sediments are usually low, however high (>15%) CaCO_3_ contents occur at shallow water depths (150–200 m) on the narrow shelves of the eastern Weddell Sea and at a depth range of 600–900 m on the broader and deeper shelves of the Amundsen, Bellingshausen and western Weddell Seas (Hauck et al., 2012).

Regions with high primary production, such as the Ross Sea and the western Antarctic Peninsula, have generally low CaCO_3_ contents in the surface sediments as carbonate produced by benthic organisms is subsequently dissolved and thus not preserved. Conversely, carbonate contents of sediments in areas of low primary productivity reflect the concentration of planktonic foraminifera that are especially abundant in sea ice (Hauck et al., 2012). As carbonate-rich sediments on the continental shelf can rapidly react to the decreasing Ω with respect to carbonate minerals (Haese et al., 2014), OA will potentially have important implications to their distribution and, consequently, the habitats of benthic organisms.

#### 1.1.3 Carbonate mineral composition of shells and skeletons

Ocean acidification will not only impact on carbonate mineral composition in surface sediments of coastal and continental shelf environments but also on diverse shelf assemblages of marine calcifiers. Globally, most scleractinian cold-water corals (>70%) will be exposed to undersaturated conditions by 2100 when the ASH is expected to reach surface waters (Guinotte et al., 2006; Turley et al., 2007). Many SO organisms may not cope with predicted rapid changes as this region is characterized by low interannual variability in surface ocean [CO_3_^2−^] (Orr et al., 2005; Conrad and Lovenduski, 2015). OA may consequently impact biomineralization of their skeletons and promote their dissolution (Andersson et al., 2008; Fabry, 2008).

While there is uncertainty in predictions about winners and losers under climate-change, the CaCO_3_ form of shells and skeletons, when combined with other biological traits, may be a key factor in predicting their response to change. Carbonate shells and skeletons may be composed of three different phases: aragonite, calcite and Mg-calcite. Calcite and aragonite are polymorphs (different mineral structure) of CaCO_3_. Mg-calcite has the same mineral structure as calcite but some Ca^2+^ ions have been replaced by Mg^2+^ ions. These CaCO_3_ minerals have different physical and chemical properties. The structure of aragonite is less stable, and thus more soluble, than that of calcite. A high percentage of marine calcifiers incorporate significant amounts of Mg into their skeletons, which also reduces mineral stability. Their shells and skeletons are often categorized as low-Mg calcite (LMC; 0–4 wt% MgCO_3_), intermediate-Mg calcite (IMC; 4–8 wt% MgCO_3_) and high-Mg calcite (HMC; >8 wt% MgCO_3_) (Rucker and Carver, 1969). Thus, HMC shells or skeletons are also more soluble than those comprised of LMC and pure calcite and even, in some situations, aragonite; consequently, they may be most susceptible to OA (Andersson et al., 2008; Fabry, 2008). However, some studies suggest that the acid-base status of a species may have more influence on their sensitivity to OA than the CaCO_3_ composition of their shell or skeleton (Collard et al., 2015; Duquette et al., 2018).

Ocean warming may exacerbate the effects of OA in species with Mg-calcite shells/ skeletons as Mg content in calcite generally increases with seawater temperature and thereby increases skeletal solubility. Ocean warming, in synergy with OA, is thus forecasted to increase the vulnerability of some marine calcifiers, particularly taxa depositing more soluble CaCO_3_ mineral phases (aragonite or HMC) (Andersson et al., 2008). However, the IMC or LMC cold-water species, potentially less soluble than those from warmer waters, may also be vulnerable to near-future OA as Antarctica is acidifying at a faster rate than much of the rest of the global ocean (Fabry et al., 2009). In addition, the species with limited thermotolerance and/or mobility (e.g. some cold-water species) are of special concern as they will not be able to shift their distribution to higher latitudes to escape from these conditions.

Information on skeletal mineralogy of Antarctic marine calcifiers is however limited to only a few taxa and particular geographic and bathymetric ranges (Borisenko and Gontar, 1991; Taylor et al., 2009; McClintock et al., 2011; Figuerola et al., 2015, 2019; Krzeminska et al., 2016). For instance, little attention has been paid to Antarctic benthic communities from the deep-sea and important areas such as the Vulnerable Marine Ecosystem (VME) in East Antarctica (Post et al., 2010). These communities are characterized by high species richness of marine calcifiers, in particular habitat-forming bryozoans and hydrocorals (Stylasteridae) (Beaman and Harris, 2005; Post et al., 2010; Bax and Cairns, 2014). Moreover, many studies have focused on the impacts of OA without considering its interactions with other environmental (e.g. warming, food quality or salinity) and biological (e.g. skeletal growth rate) factors which are involved in mediating the susceptibility of marine calcifiers to OA, for example, through controlling the incorporation of Mg into the skeletal calcite (Chave, 1954; Andersson et al., 2008; Hermans et al., 2010; Sewell and Hofmann, 2011; Swezey et al., 2017). Here we combine a review and a quantitative meta-analysis to provide an overview of the current state of knowledge about skeletal mineralogy of major taxonomic groups of SO marine calcifiers and to make predictions about how OA may affect different taxa.

## 2 Methods

In order to predict how OA will affect common marine calcifiers from the SO, environmental and biological data for a broad range of SO marine calcifiers were compiled. These included species geographic range, skeletal mineralogy, the calcification responses and strategies to overcome OA. Their CaCO_3_ composition was reported as aragonite and/or calcite. Calcite composition was further classified as LMC, IMC and HMC (Rucker and Carver, 1969). Data previously reported as mol% MgCO_3_ were converted to weight % (wt%) MgCO_3_ for comparisons. Individual SO maps with species distributions were then produced for two groups (echinoderms and bryozoans) with Mg-calcite skeletons for which there is relatively good data and for benthic species identified here as particularly vulnerable to ocean acidification.

With the aim of performing a meta-analysis, database searches were conducted to compile all peer-reviewed journal articles and literature reviews that investigated the effect of altered seawater carbonate chemistry on SO marine calcifiers using the criteria and methods outlined in Hancock et al. (2020) (database searches completed May 2020). Articles were screened to include studies that manipulated the carbonate chemistry to investigate the effects of OA on specifically SO marine calcifiers, and after manual screening, 20 studies remained for inclusion in the meta-analysis (Fig. S1). Data included bivalves (Mollusca; Bivalvia), brachiopods (Brachiopoda), limpets (Mollusca; Gastropoda; Patellogastropoda), pteropods (Mollusca; Gastropoda; Pteropoda), sea snails (Mollusca; Gastropoda; Trochida), sea stars (Echinodermata; Asteroidea), sea urchins (Echinodermata; Echinoidea) and coccolithorphores (Haptophyta; Coccolithophyceae). The effects of OA on the following biological responses were investigated: development (bivalves, sea stars and sea urchins; fertilization rate or normal development), growth rate (brachiopods, pteropods and coccolithophores), shell state (bivalves, limpets, pteropods and sea snails; adult calcification or dissolution rate), coccolith volume (coccolithophores) and survival (sea urchins). The statistical analyses of the data are explained in detail in Hancock et al. (2020). In brief, for each study the ln-transformed response ratio to OA was calculated using the ambient treatment (<500 μatm) and highest CO_2_/lowest pH treatment (up to μatm 1500 CO_2_). For all response ratios, a positive ratio is a positive response to OA, a negative response ratio is a negative response to OA, and a response ratio of zero indicates there is no effect of OA. These response ratios were then weighted by the variance, where studies with a lower standard error and higher sample size were weighted more heavily than those with a higher standard error and lower sample size (Hedges and Olkin, 1985). Meta-analyses were conducted with all studies combined and separated based on the mineralogical composition of the study organism (as outline above).

## 3 Results

For the Mg calcite-producing species, the calcite composition was classified in a total of 123 species belonging to 89 bryozoan and 34 echinoderm species (Tables S1-S2). Individual SO maps showed a general pattern of increase of aragonite and the Mg content in calcite towards lower latitudes in bryozoans and echinoderms (Fig. 2).

**Fig. 2.**
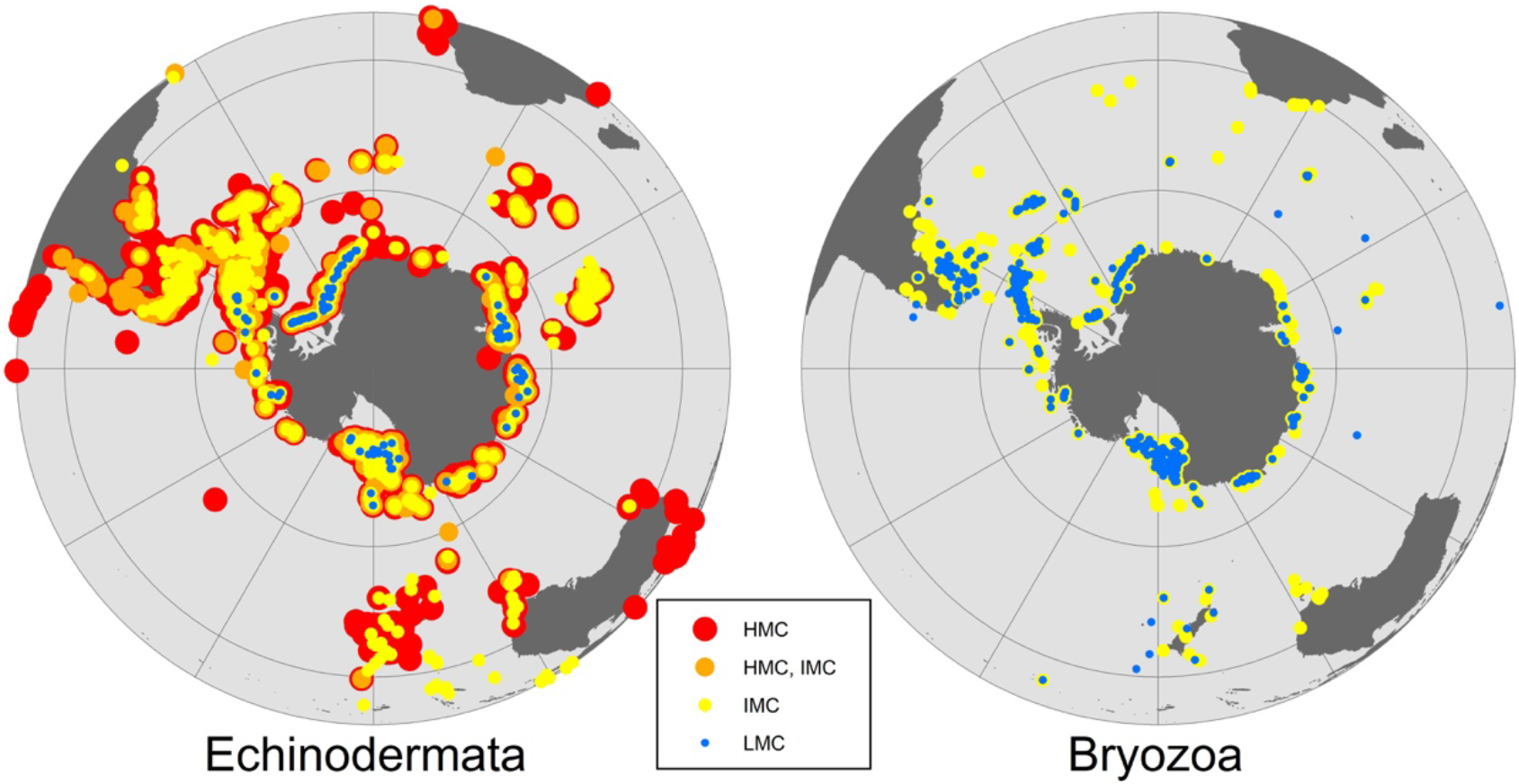
General distribution patterns for Echinodermata (24 HMC, 3 HMC/IMC, 6 IMC and 1 LMC species) and Bryozoa (59 IMC and 30 LMC species) based on Mg-calcite distribution in the Southern Ocean.

Twenty studies on the effect of altered carbonate chemistry on marine calcifiers south of 60°S were included in the meta-analysis (Fig. S1 and Table S3). A total of 25 response ratios (two for bivalves, two for brachiopods, four for gastropods, two for sea stars, nine for sea urchins, two for pteropods, one for sea snail, one for patellogastropoda and six for coccolithophores) were calculated from the 20 studies (some studies reported more than one experiment and/or species response). With all response ratios combined, there was a negative response to OA in marine calcifiers (Fig. 3). However, there was significant heterogeneity (Q = 48669.264, df = 24, p < 0.001; Table S3) indicating that the variability between the studies is greater than expected by chance alone. The response varied depending on the mineralogical composition of the organism (Fig. 3). While there was a negative response to OA in calcitic, aragonitic and HMC species, LMC species were mostly unaffected. Significant heterogeneity was observed for aragonite, calcite and HMC (p < 0.001) but not for LMC (Q = 0.6085, df = 1, p = 0.4354) (Table S3). The echinoderms, *Sterechinus neumayeri* and *Odontaster validus*, were sensitive to increased CO_2_ above current levels (>500 μatm), with a negative effect on their development. The shell state and growth of the pteropod *Limacina helicina* and the shell state of the sea snail *Margarella antarctica* were negatively affected at moderate CO_2_ levels (>800 μatm). The response was also negative in CO_2_ treatments exceeding 1,000 μatm in the brachiopod *Liothyrella uva* (growth), the bivalve *Laternula ellitica* (development and shell state), the coccolithophore *Emiliania huxleyi* (growth) and the sea urchin *S. neumayeri* (survival) (see Fig. S1 for details).

**Fig. 3.**
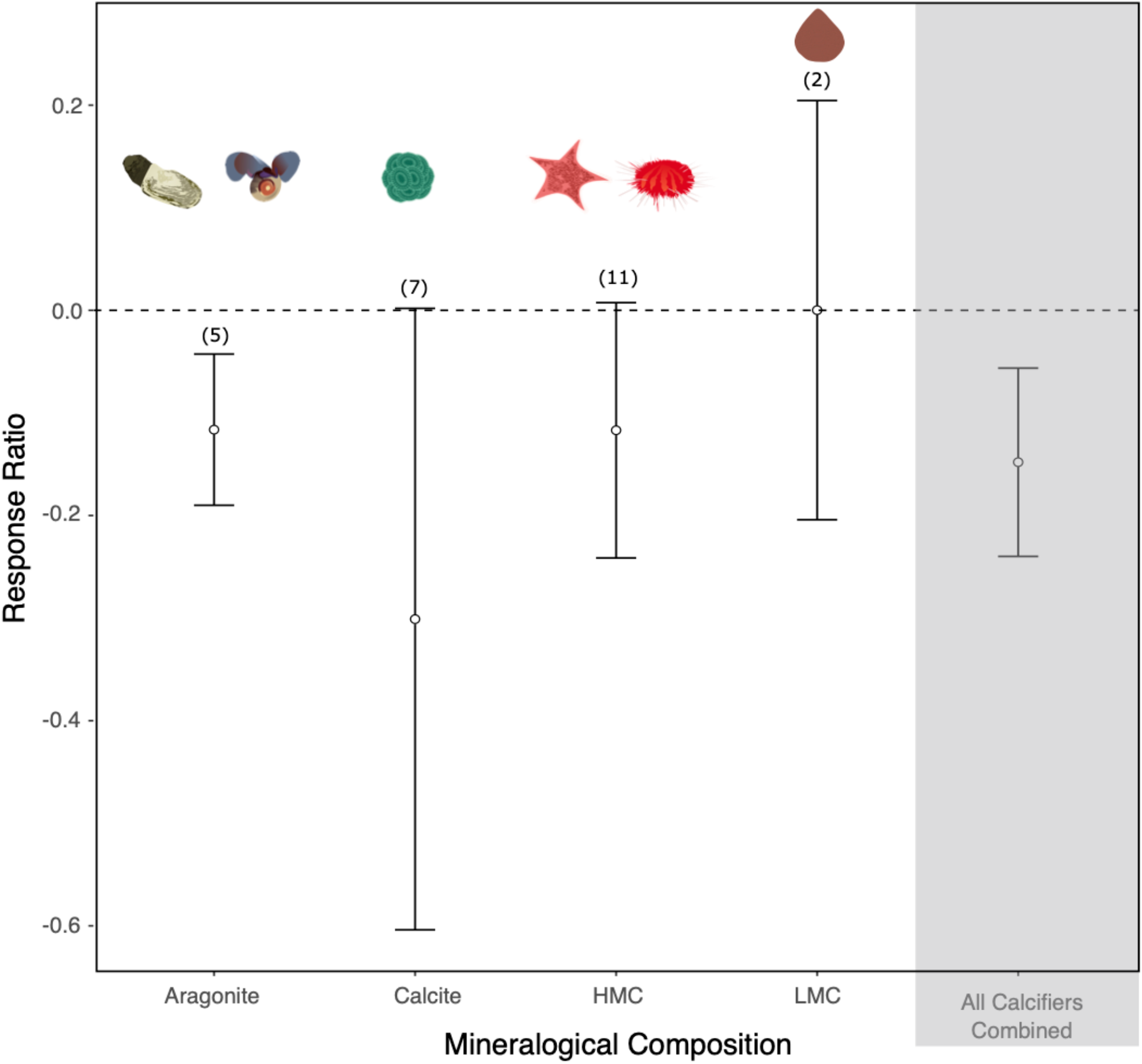
The effect of ocean acidification on aragonitic, calcitic, high-Mg calcite (HMC) and low-Mg calcite (LMC) species. Mean response ratios and 95% confidence intervals are shown, with the number of data points in each category given in brackets. A mean response ratio of zero (hashed line) indicates no effect. Background shading indicates response ratios of all the phyla.

Information on the vulnerability of a range of different common Antarctic phyla was assessed based on their mineralogy-related sensitivity and their strategies to overcome OA (Table 1).

**Table 1.**
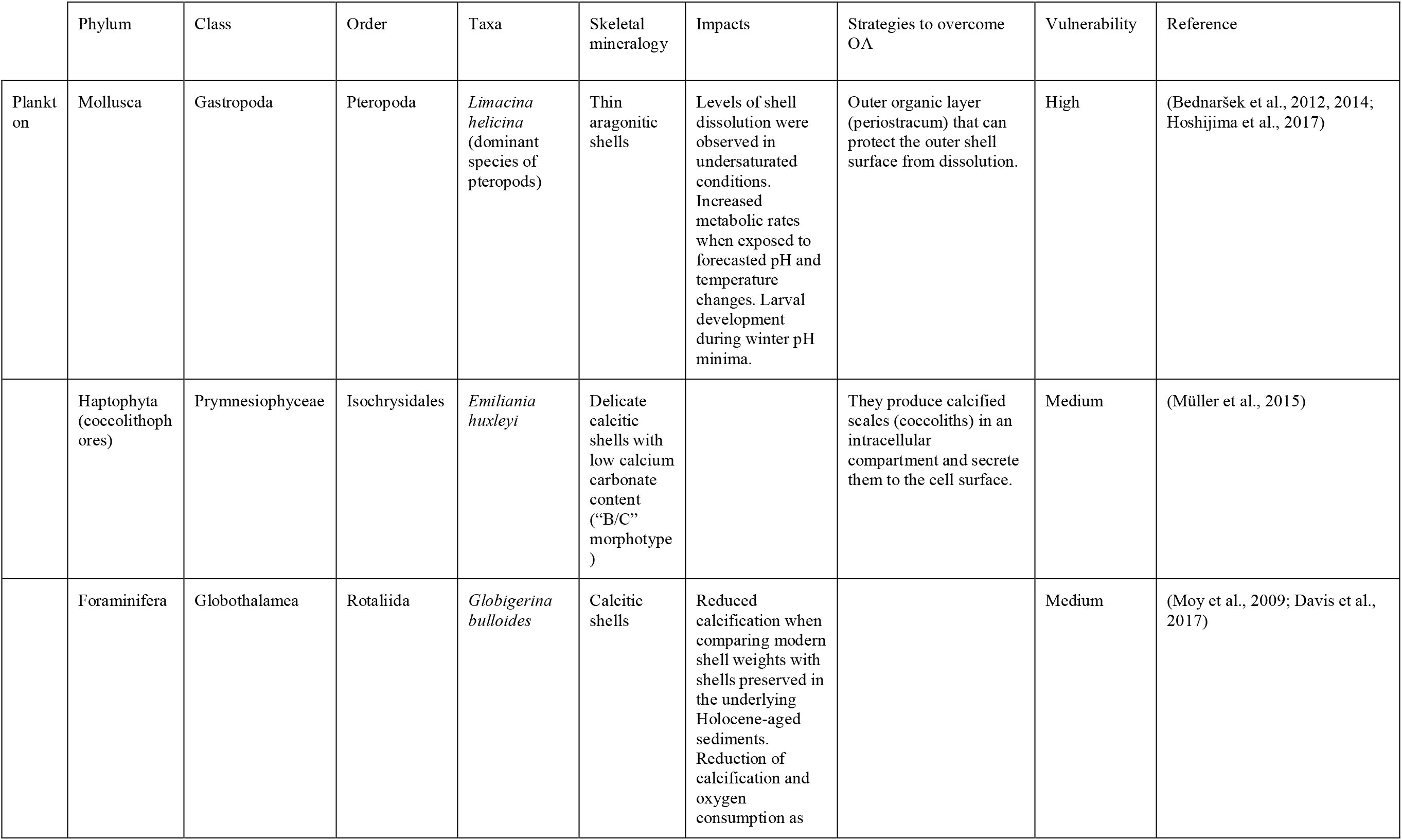
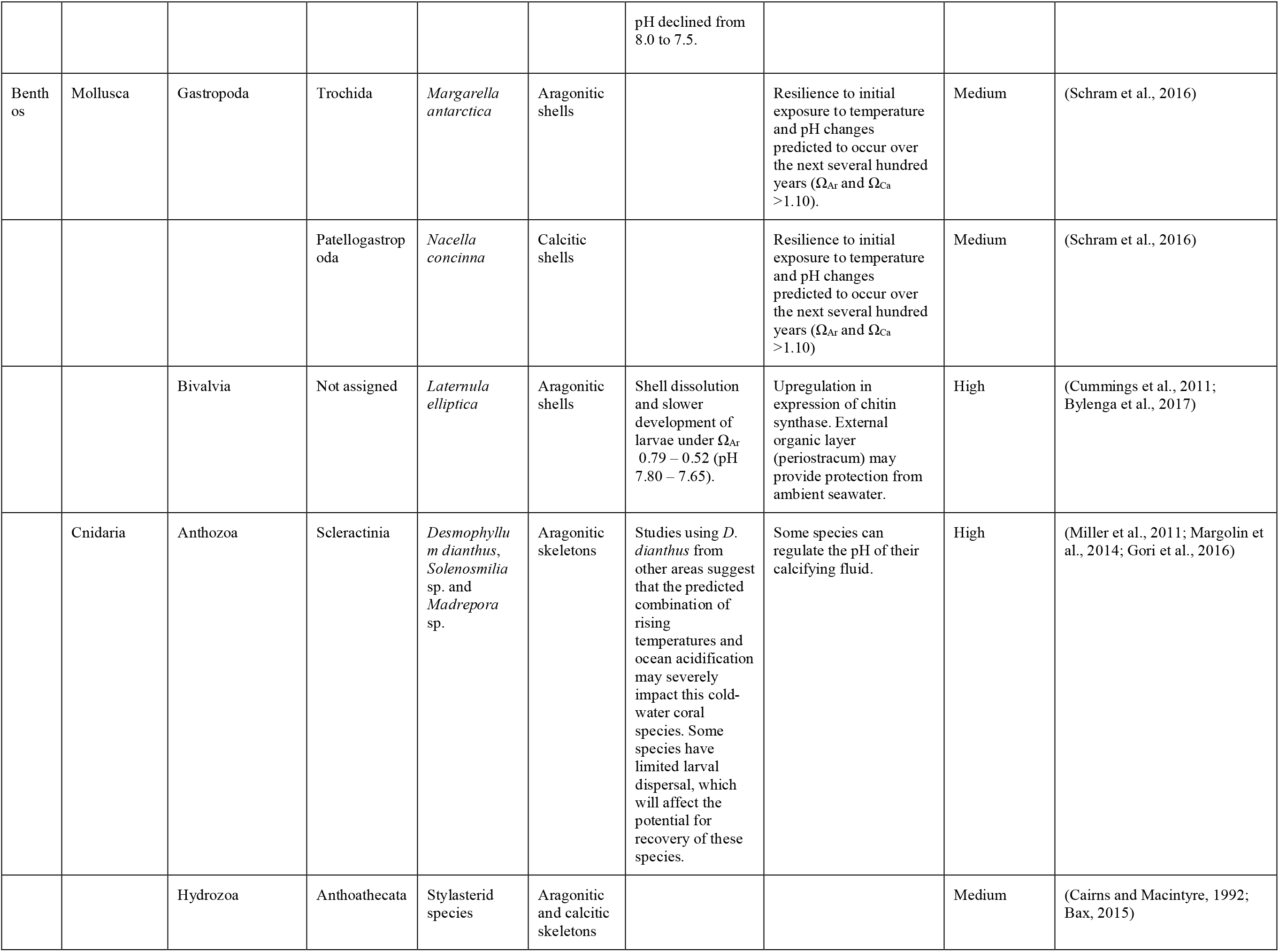
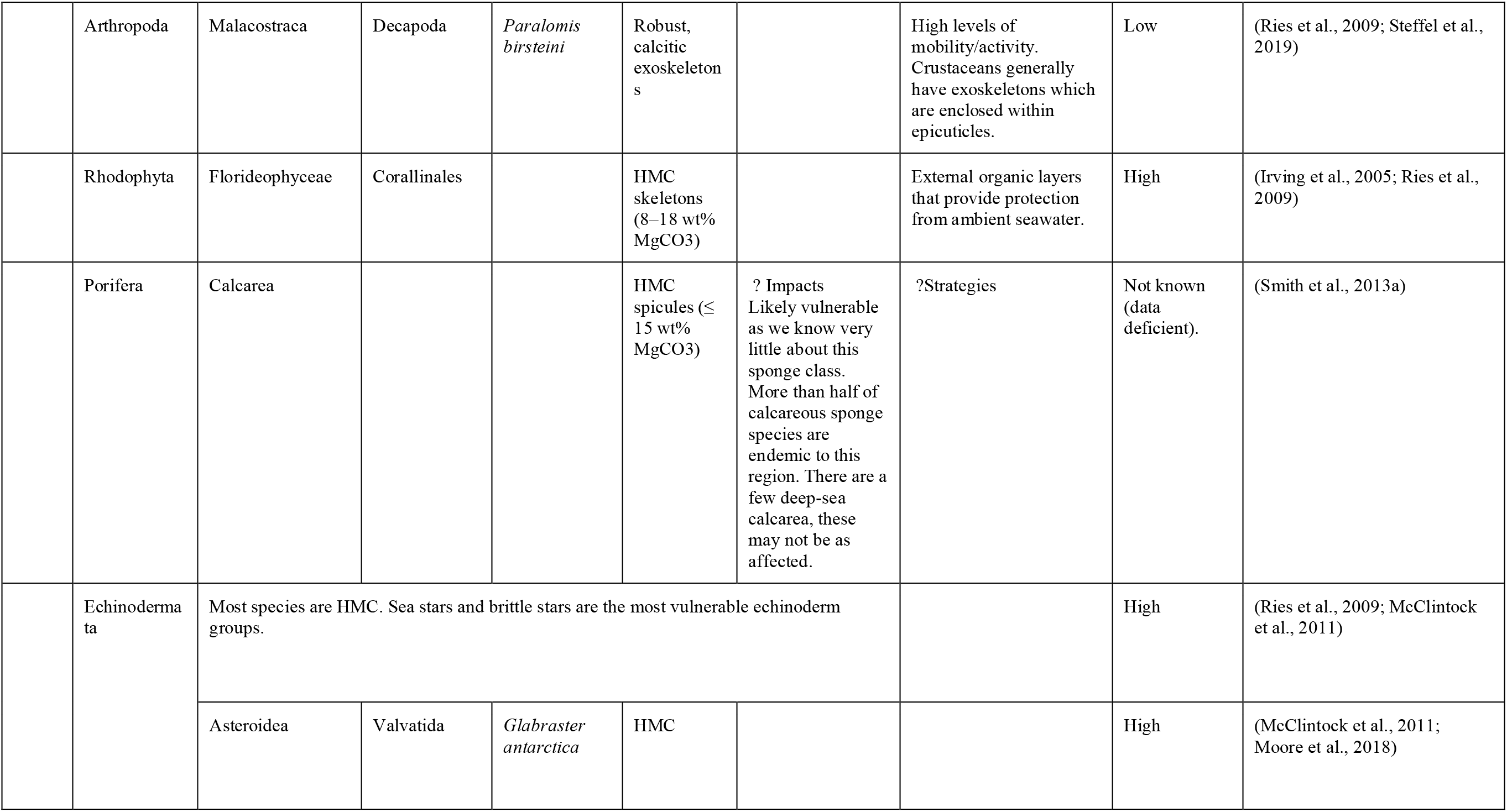
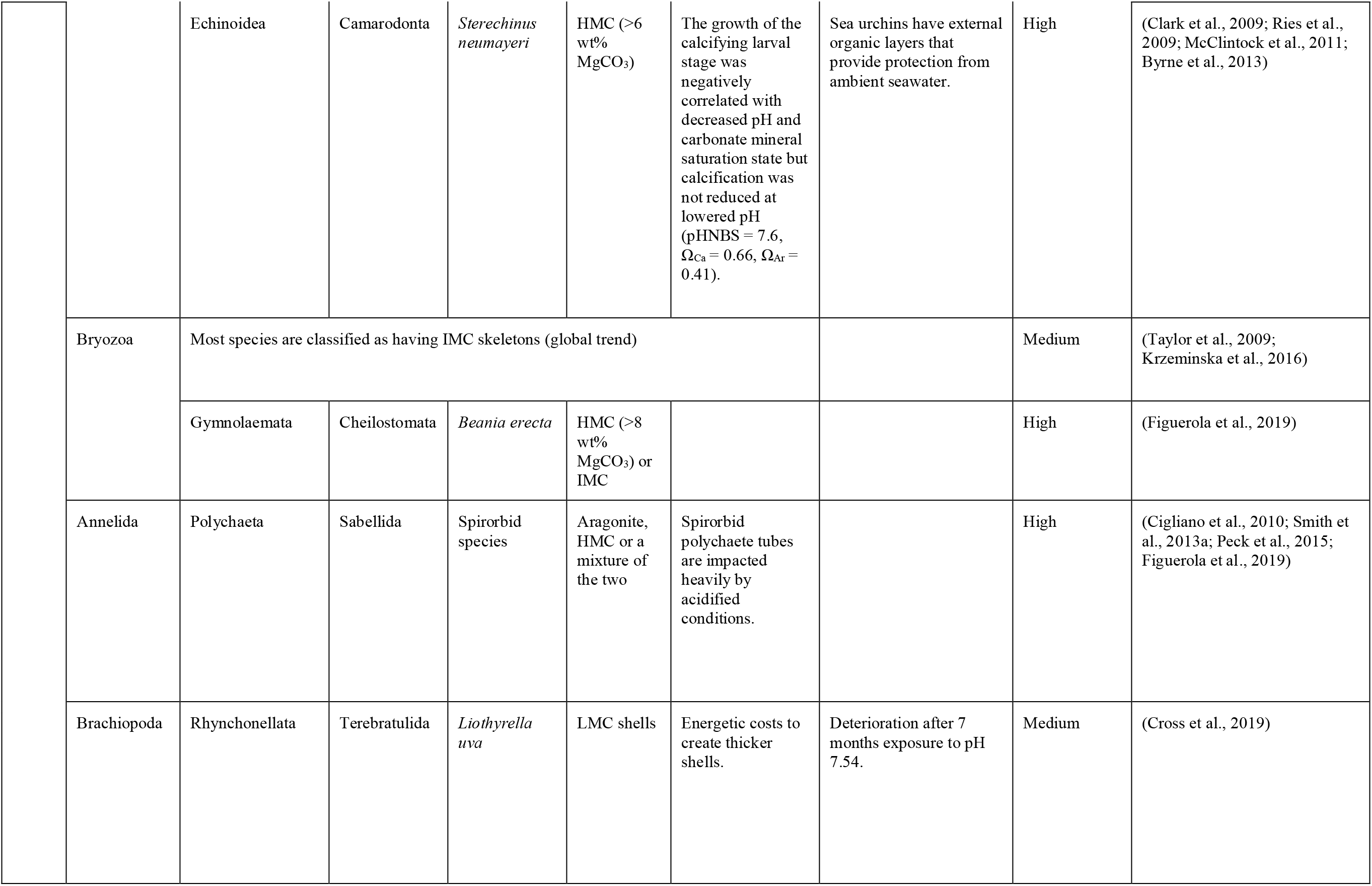
Potential vulnerability of a range of different common Antarctic phyla based on their mineralogy-related sensitivity and their strategies to overcome OA. LMC: low-Mg calcite (0–4 wt% MgCO_3_); IMC: intermediate Mg calcite (4–8 wt% MgCO_3_); HMC: high Mg calcite (> 8 wt% MgCO_3_).

## 4 Mineralogy-related sensitivity

Several common Antarctic marine calcifiers secrete more soluble CaCO_3_ mineral phases: aragonite (e.g. scleractinian cold-water corals, infaunal bivalves, pteropods) or HMC (e.g. bryozoans and echinoderms) (Turley et al., 2007; McClintock et al., 2009, 2011; Figuerola et al., 2019). These groups play important roles in ecosystem functioning acting as a food source of carnivorous zooplankton, fishes and other predators (e.g. pteropods), creating habitats used as spawning, nursery and feeding areas of higher trophic levels (e.g. corals, spirorbid polychaetes), structuring benthic communities as a result of grazing/predation (e.g. echinoderms) (Dayton et al., 1974; Prather et al., 2013; Wright and Gribben, 2017) and/or contributing to marine carbon cycling and storage (e.g. foraminifera, coccolithophores, molluscs) (Lebrato et al., 2010). Moreover, some Antarctic calcifying species belonging to these groups are characterized by circumpolar distributions and broad bathymetric ranges, and may make potentially good indicator species of chemical change in the SO (Dayton et al., 1974; Brandt et al., 2007; Barnes and Kuklinski, 2010; Figuerola et al., 2012). Under expected ASH shoaling, these taxa could find refuge in shallower waters. Yet, more molecular studies are needed as eurybathy may be less prevalent than once thought, with the recent increase in the detection of cryptic species (Baird et al., 2011; Allcock and Strugnell, 2012).

### 4.1 Aragonite and/or calcite-producing species

The anticipated shallowing of the ASH and associated aragonite undersaturation in shallow waters is of special concern for benthic molluscs and zooplankton species such as pteropods which build aragonitic shells and/or skeletons. SO pteropods are the major pelagic producers of aragonite and comprise up to one-quarter of total zooplankton biomass in several Antarctic waters (the Ross Sea, Weddell Sea and East Antarctica) with *Limacina helicina* (Phipps, 1774) as the dominant species (McNeil and Matear, 2008). These free-swimming marine gastropods are considered sentinels of OA as their thin aragonitic shells are vulnerable to OA (Bednaršek et al., 2014). Levels of shell dissolution were observed when individuals of *L. helicina antarctica* were exposed to undersaturated conditions (ΩA = 0.8) (Bednaršek et al., 2012, 2014). Its metabolic rate also increased when exposed to lowered pH (pH 7.7) and a high temperature (4°C) likely related to increased costs for shell repair (Hoshijima et al., 2017). *L. helicina* has a life cycle of 1–2 years and its veliger larva develops during winter, when the expected early aragonite undersaturation will occur. If *L. helicina* is not able to cope with anticipated OA, it will likely lead to cascading impacts through higher trophic levels (Seibel and Dierssen, 2003). Similarly, the D-larval stage of the common Antarctic bivalve *Laternula elliptica* (P. P. King, 1832), a large bivalve with a circumpolar distribution (Fig. 4), showed shell dissolution and slower development under Ω_Ar_ 0.79 – 0.52 (pH 7.80 – 7.65) (Bylenga et al., 2015, 2017). These results are expected as their larvae possess aragonitic shells. Empty adult shells of other SO molluscs (bivalves and limpets), including calcitic species, also suffered significant dissolution when exposed to reduced pH (pH = 7.4, Ω_Ar_ = 0.47, Ω_Ca_ = 0.74) (McClintock et al., 2009). In contrast, net calcification in individuals of the limpet *Nacella concinna* (Strebel, 1908) and the snail *Margarella antarctica* (Lamy, 1906), with calcitic (Mg-calcite; J.B. Schram, unpublished X-ray diffraction data) and aragonitic shells, respectively, was resilient to initial exposure to temperature and pH changes predicted to occur over the next several hundred years (Ω_Ar_ and Ω_Ca_ >1.10) (Schram et al., 2016).

**Fig. 4.**
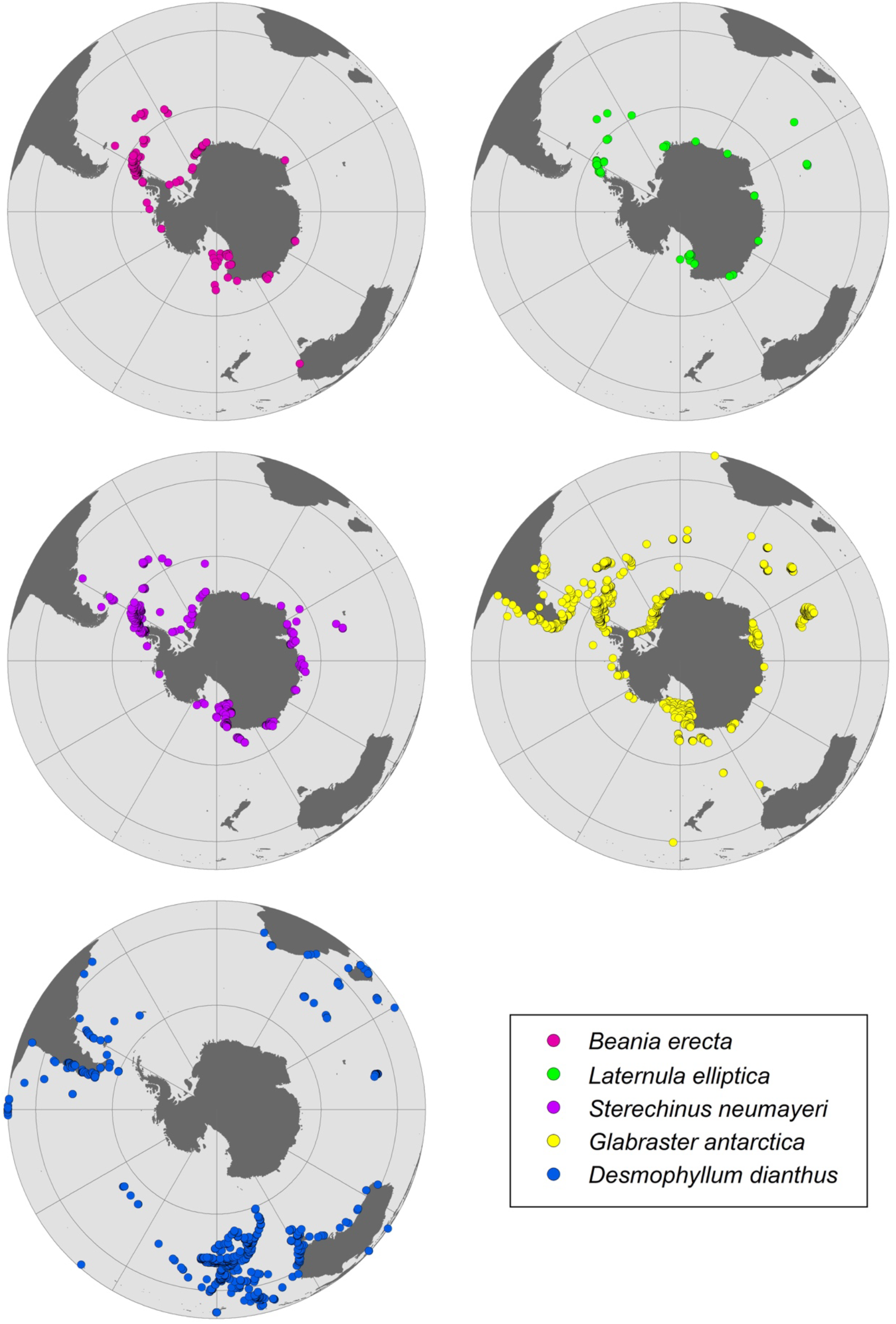
Distribution patterns for benthic species from the Southern Ocean identified here as particularly vulnerable to ocean acidification.

Coccolithophores, globally ubiquitous, are the major phytoplanktonic calcifier group, accounting for nearly half of global CaCO_3_ production (Moheimani et al., 2012). These unicellular flagellate algae play a significant role in the SO carbon cycle, contributing 17 % to annually integrated net primary productivity south of 30° S (Nissen et al., 2018). A recent study, including coccolithophores, showed there were taxon-specific and local variation in phytoplankton community responses to increased pCO_2_ and light intensity (Donahue et al., 2019). *Emiliania huxleyi* (Lohmann) W.W.Hay & H.P.Mohler, 1967, the most abundant species, has different morphotypes that exhibit a clear north-to-south distribution gradient, with the SO “B/C” morphotype forming delicate coccoliths with relatively low CaCO_3_ (calcite) content compared to the other more calcified morphotypes from lower latitudes (Müller et al., 2015). However, the rare occurrence of heavily calcified morphotypes in higher latitudes/lower pH waters highlights that coccolithophores can calcify in lower CaCO_3_ conditions (Beaufort et al., 2011). Three morphotypes from different latitudes reduced their rate of calcification under simulated OA scenarios; the B/C morphotype the most sensitive, although even in the more resilient morphotype A a three-fold reduction in calcification was observed (Müller et al., 2015, 2017). However, an evolutionary study shows that calcification can be partly restored under OA conditions over approximately one year or 500-generations (Lohbeck et al., 2012). Given the shorter turn-over time of phytoplankton compared to higher-level organisms, the ability of coccolithophores to rapidly adapt to changing environmental conditions could help maintain their functionality within the ecosystem.

Globally, gorgonian corals (soft corals) account for the majority of OA coral literature (McFadden et al., 2010; Thresher et al., 2010; Watling et al., 2011), and data beyond presence/absence and taxonomic data is available for some widely distributed Antarctic species (Taylor and Rogers, 2015; Moore et al., 2017). Comparatively, there is very little data on Antarctic coral species beyond taxonomic descriptions (Cairns, 1982, 1983), and biogeographic studies (Bax and Cairns, 2014), leaving their calcification responses to OA largely unknown, despite particular vulnerability (Tittensor et al., 2010). The only exception is the cosmopolitan aragonite scleractinian coral *Desmophyllum dianthus* (Fig. 4) (Esper, 1794), for which a reasonable literature relating to OA exists (Miller et al., 2011; Fillinger and Richter, 2013; Jantzen et al., 2013). However, most of these studies are based on populations outside of Antarctica, predominantly within the Chilean fjords. Globally, gorgonian corals account for the majority of OA coral literature (McFadden et al., 2010; Thresher et al., 2010; Watling et al., 2011). Cold-water corals form a dominant component of VMEs throughout the Antarctic and sub-Antarctic, including gorgonian corals (Moore et al., 2017), the colonial scleractinian corals, such as *Solenosmilia* and *Madrepora* (Cairns, 1982; Cairns and Polonio, 2013) and stylasterid corals (e.g. *Errina* sp.) (Bax and Cairns, 2014). These coral groups have been shown to have differential carbonate mineralogy: gorgonians: calcite/aragonite (Thresher et al., 2011); scleractinians: aragonite (Margolin et al., 2014), and stylasterids: calcite/aragonite (Cairns and Macintyre, 1992). Therefore, they may be differentially affected by OA. Cold-water corals are also generally exposed to relatively low levels of environmental variability, thus are likely to be more sensitive to rapid changes in seawater chemistry (Thresher et al., 2011). The discovery of large field-like aggregations of deep-sea stylasterid coral reefs in the Antarctic benthos and their classification as a VME under the Commission for the Conservation of Antarctic Marine Living Resources (CCAMLR) highlights the conservation importance, and recognises the role these reefs play in the maintenance of biodiversity (Bax and Cairns, 2014). Notably, stylasterid corals have the ability to utilise both aragonite and calcite (Cairns and Macintyre, 1992), and may have a greater capacity to acclimate to changing oceanic pH than purely aragonite calcifiers (Bax and Cairns, 2014). Therefore, there is a strong need to conserve and better understand the role they may play in the maintenance of Antarctic biodiversity under predicted climate change scenarios (Bax et al., in review, this issue).

### 4.2 Mg calcite-producing species

High latitude surface seawater is currently undersaturated with respect to Mg-calcite minerals containing >10 wt% MgCO_3_ (Andersson et al., 2008). The skeletal Mg-calcite composition has been determined for a considerable proportion of species from several Antarctic taxa (e.g. echinoderms and bryozoans) although there is still a gap in knowledge of other taxa and geographic variation in general.

Most foraminifera genera have calcitic shells with various amounts of Mg, ranging from LMC to HMC (Blackmon and Todd, 1959). Planktonic foraminifera are ubiquitous in the SO and a major component of calcifiers among marine zooplankton, with ∼25% settling on the seafloor and their calcitic shells contributing ∼32–80% to the total deep-marine calcite budget (Schiebel, 2002). It is known that reductions in pH and [CO_3_^2−^] generally alter the performance of this group, together with pteropods and coccolithophores (Barker and Elderfield, 2002; Manno et al., 2012). *Globigerina bulloides* d’Orbigny, 1826, one of the most commonly used species for paleoreconstructions in high latitudes and found from tropical to cold waters, alternates high and low Mg/Ca layers embedded within its test (Anand and Elderfield, 2005). The common SO species have shown reduced calcification when comparing shells preserved in the underlying Holocene-aged sediments with modern shell weights (Moy et al., 2009). The authors linked the change to anthropogenic OA. Other lab and field studies using different species of planktonic foraminifera showed the same trend (Manno et al., 2012; Marshall et al., 2013). Consistent with these previous findings, a recent study showed a reduction of calcification and oxygen consumption of *G. bulloides* as pH declined from 8.0 to 7.5 (Davis et al., 2017). The lack of studies on the OA effects on foraminifera in the SO, and in general, highlights the importance of more research on this field to make more accurate prediction.

The tissue of crustose coralline algae skeletons typically ranges from 8–18 wt% MgCO_3_ (Chave, 1954), and they may have the potential to act as ‘sentinel’ species. The distribution of crustose coralline algae in Antarctica is not well known and the taxonomy is not well resolved either (Wiencke et al., 2014). Surprisingly, this group is abundant and diverse worldwide and in a range of habitats on hard substrata of the continental shelves. It has been found to cover up to 70–80% of the substrate under macroalgal canopies in Antarctica in shallow water (Irving et al., 2005). Crustose coralline algae may also be a key substrate for benthos in some Antarctic fjords at 10–30 m (Wiencke et al., 2014) and it has colonised most newly ice-free areas (Quartino et al., 2013). There is one main species, *Phymatolithon foecundum* L. Düwel & S. Wegeberg (Alongi et al., 2002), which has been reported to depths of 70 m (Cormaci et al., 2000) in the Ross Sea. While the mineralogy of this species in the Antarctic has not been investigated, a recent investigation of four *Phymatolithon* spp. in the north Atlantic revealed that, within a single crust, Mg content of the carbonate ranged from 8 to 20 mol% MgCO_3_ (Nash and Adey, 2017). Furthermore, the proportions of carbonate types varied within the tissues, and also between species (Nash and Adey, 2017), making it difficult to predict OA impacts. Antarctic crustose macroalgae may be resilient to OA, with a study conducted on *Clathromorphum obtectulum* (Foslie) W.H.Adey, 1970 observing no negative impacts on CaCO_3_ content and Mg/Ca ratio with OA (Schoenrock et al., 2016).

Sponges are often species-rich, dominant, high biomass, habitat-providers in many Antarctic seafloors, with calcareous sponges representing more than 12% of all species, which is far higher than the global average (Soest et al., 2012). Calcareous sponges are also found to be highly endemic, and have two genera found only in this region (Downey et al., 2012). Species belonging to the class Calcarea generally produce CaCO_3_ spicules with very high Mg contents (**≤**15 wt% MgCO_3_), and are likely to be vulnerable to projected lowered pH (Smith et al., 2013a). Close to 90% of calcareous sponges are only found on the shelves of the Antarctic continent and sub-Antarctic continents (Janussen and Downey, 2014), and most of these species have very narrow longitudinal and latitudinal ranges, which likely increases their vulnerability to predicted OA in the SO.

Recent studies have investigated the skeletal Mg contents in calcite of 29 Antarctic echinoderm species to make predictions about how OA may affect different groups (McClintock et al., 2011; Duquette et al., 2018) (Table S1). Most species were HMC although variation of skeletal Mg-calcite existed between taxonomic classes (asteroids, ophiuroids, echinoids, and crinoids), and for the same species from different locations. McClintock et al. (2011) suggested sea stars (e.g. *Glabraster antarctica* (E. A. Smith, 1876)), which are Antarctic keystone predators (Dayton et al., 1974), and brittle stars, with higher mean Mg values, will probably be the first echinoderms to be affected by OA. The fertilization and early development of the common Antarctic sea urchin *Sterechinus neumayeri* (Meissner, 1900) were not affected by near-future changes and its fertilization success only declined significantly when exposed to one stressor (pH) at reduced levels predicted for 2300 (decreases of ~ 0.7 pH units) (Ericson et al., 2010; Ho et al., 2013). However, a negative interactive effect of projected changes in seawater temperature (3°C) and pH was found on the percentage of fertilization (11% reduction at pH 7.5) (Ericson et al., 2012) and on the cleavage success (pH 7.6) (Foo et al., 2016) although the results may vary depending on sperm concentrations, population or experimental conditions (Sewell et al., 2014). The growth of the calcifying larval stage of this species was also found to be negatively correlated with decreased pH and Ω (Byrne et al., 2013) but calcification was not reduced at lowered pH (pH_NBS_ = 7.6, Ω_Ca_ = 0.66, Ω_Ar_ = 0.41)(Clark et al., 2009). Also, its larvae had shorter arm lengths when exposed to undersaturated conditions (730 μatm, Ω=0.82) (Yu et al., 2013). Similarly, normal development of the seastar *Odontaster validus* Koehler, 1906 was reduced at low pH (7.8) (Karelitz et al., 2017) as was survival and growth of the larvae at pH_NIST_ 7.6 (Ω_Ca_ = 0.89, Ω_Ar_ = 0.57) (Gonzalez-Bernat et al., 2013), indicating that they are unable to adjust to changes in extracellular pH, even though Asteroidea are non-calcifying during their larval development. The latter study illustrates how polar organisms are affected by characteristics of OA other than direct effects on carbonate structures, and which likely explain their different responses and sensitivities and lack of relationship to Ω (Ries et al., 2009; Waldbusser et al., 2015).

The Mg content from individuals of approximately 90 Antarctic bryozoan species (nearly one-quarter of all Antarctic bryozoan species) demonstrates extreme intraspecific and interspecific variation (Borisenko and Gontar, 1991; Taylor et al., 2009; Loxton et al., 2013, 2014; Figuerola et al., 2015, 2019; Krzeminska et al., 2016) (Table S2). These studies showed most species consisted of ICM. Taylor et al. (2009) previously confirmed the absence of aragonitic bryozoan species at high latitudes (>40°S) and the rarity of bimineralic species. Figuerola and coauthors (2019) found the circumpolar cheilostome *Beania erecta* Waters, 1904 (Fig. 4) may be particularly vulnerable to global ocean surface pH reductions of 0.3–0.5 units by the year 2100 as some specimens had HMC skeletons. The Antarctic cyclostome *Fasciculipora ramosa* d’Orbigny, 1842 also seems to be influenced to some degree by environmental factors, as shown by the significant variability in branch diameter and Mg levels among depths (Figuerola et al., 2015, 2017).

Encrusting biofouling communities like spirorbid polychaetes and bryozoans are amongst the most remarkable early Antarctic colonizers (Stark, 2008). Spirorbid polychaetes, with r-selected life-history strategies, are impacted heavily by acidified conditions as their tubes are mainly composed of aragonite, HMC or a mixture of the two (Cigliano et al., 2010; Smith et al., 2013b; Peck et al., 2015). A recent study did not find any temporal variation over 6 yrs in the wt% MgCO_3_ in calcite in Antarctic bryozoans and spirorbid polychaetes, suggesting these particular taxa may be a good indicator species for the potential effects of environmental change (Figuerola et al., 2019).

Brachiopods have some representatives in Antarctic waters. In particular, the rhynchonelliform brachiopods are locally important members in shallow water communities and have LMC shells (Barnes and Peck, 1996). The circumpolar rhynchonelliform brachiopod *Liothyrella uva* (Broderip, 1833) showed deterioration after 7 months exposure to pH 7.54 (Ω_Ca_ = 0.5, Ω_Ar_ = 0.3; (Cross et al., 2019) although the predicted acidified conditions did not alter shell growth rates and the ability to shell repair (Cross et al., 2014).

In crustaceans, exoskeletons of several species have also been shown to contain Mg-calcite, although they generally have a very complex mineralogy (combination of calcite, Mg-calcite, calcium phosphate and chitin) compared to skeletons of other groups like bryozoans (Chave, 1954; Andersson and Mackenzie, 2011). King crabs (Lithodidae), inhabiting deep water habitats of the SO that are already undersaturated in aragonite (Steffel et al., 2019), produce calcareous, even robust, exoskeletons (e.g. *Paralomis birsteini* Macpherson, 1988). As projected global warming removes thermal barriers controlling their distribution, king crabs may expand their bathymetric range to the Antarctic continental shelf, a region currently characterized by the absence of durophagous (skeleton-breaking) predators (Griffiths et al., 2013). Therefore, a range of calcifying invertebrates (e.g. echinoderms and molluscs) with thin and fragile shells and skeletons would be at real risk of predation were such predators to establish on the Antarctic shelf.

Changes in seawater temperature, which strongly influences growth rate and also Ω, are also thought to drive changes in skeletal Mg content, although other environmental factors such as salinity may be influential (Chave, 1954; Mackenzie et al., 1983). There is a general trend of increase in aragonite and in Mg content in calcite towards lower latitudes in different marine calcifying taxa (Lowenstam, 1954; Taylor et al., 2009). A similar trend was found in different life stages (e.g. juveniles and larvae vs. adults) and nine skeletal elements (e.g. spines and tooth) of echinoids (Smith et al., 2016). This general pattern was also observed in individuals of echinoderms and bryozoans with LMC skeletons only reported in Antarctica and in sub-Antarctic and Antarctic regions, respectively (Fig. 2).

## 5 Strategies to overcome ocean acidification

Maintaining calcified skeletons, especially those with high Mg content, is more energetically demanding in cold waters, as the solubility of CaCO_3_ increases with decreasing temperature (Weyl, 1959). This is supported by a clear signal of decreasing skeletal investment with latitude and decreasing temperature, although a significant flexibility in some Antarctic species exists, e.g. laternulid clams have thicker shells than lower latitude congenerics (Watson et al., 2012). Under a global warming scenario, a general increase in skeletal Mg content is expected, likely making the skeletons even more vulnerable to dissolution (Chave, 1954; Andersson et al., 2008; Hermans et al., 2010; Sewell and Hofmann, 2011). Hence, energetic costs to counteract dissolution will likely have implications for many taxa.

Our meta-analysis investigating the effects of the OA on a range of biological responses (development, growth rate, survival and/or shell state) of SO calcifiers showed calcitic, aragonitic and HMC species are vulnerable to OA whereas LMC species were unaffected (Table S3), which could thus indicate a resilience to OA. A previous comprehensive meta-analysis also showed reductions in survival, calcification, growth and development of a range of marine calcifiers worldwide in response to OA (Kroeker et al., 2013). This study incorporated a data set from a previous meta-analysis, which found marine calcifiers were more commonly affected by OA than non-calcifying taxa except for crustaceans, which showed resilience (Kroeker et al., 2010). These authors concluded that the organisms depositing high Mg-calcite could be more resilient to OA than those depositing calcite or aragonite, bringing into question the use of that simplified mineralogical composition in predicting organism sensitivities to OA (Kroeker et al., 2010). A subsequent constructive debate arose, leading to several suggestions to improve the accuracy of predictions, such as including other biological variables as well as the mineralogical composition, redefining the HMC category as skeletons containing >12 mol% MgCO_3_ and excluding those with very complex mineralogy (e.g. crustaceans) (Andersson and Mackenzie, 2011; Kroeker et al., 2011). The authors also reinforced the need for more mineralogical studies to characterize a wider range of species. Further, a recent meta-analysis specifically investigated the effect of altered carbonate chemistry on marine organisms south of 60°S from 60 studies (Hancock et al., 2020). Consistent with the previous findings (discussed above), reduced survival and shell calcification rates, and increased dissolution with OA was found in adult calcifying invertebrates, along with a reduction in embryonic or larval development were also shown (Hancock et al., 2020).

Here, a compilation of the available research, including the mineralogy-related sensitivity of different marine calcifier groups, suggests that molluscs are one of the groups most sensitive to OA and that crustaceans may be more resilient (Table 2). Mechanisms likely to provide marine calcifiers with greater resilience to OA include high levels of mobility/activity and consequently higher metabolic rates (e.g. crustaceans), greater larval dispersal, capacity to regulate the pH of their calcifying fluid; and protection of shells or skeletons from the surrounding water (protective external organic layers) (Melzner et al., 2009; Ries et al., 2009). Responses and mechanisms for adaptation (permanent evolutionary modifications induced by response to repeated stressors) or acclimation (potential for an organism to adjust to changes in an environment) to the forecast changes are anticipated to be species-specific due to the phylogenetic control on the calcification process and on the Mg content in calcite in some taxa (Chave, 1954; Cairns and Macintyre, 1992).

**Table 2.** Effects of acidification on survival, calcification, growth, development, photosynthesis in marine calcifier groups from Kroeker et al. 2013, including new data of mineralogy-related sensitivity and vulnerability. Effects represented as mean percent (−) decrease in a given response, no effects (or too few studies) represented as 0, and untested variables by blank space. Groups are scored from low to high vulnerability to ocean acidification, considering the result of each response variable and skeletal mineralogy.

| Group | Survival | Calcification | Growth | Development | Photosynthesis | Mineralogy | Vulnerability |
| --- | --- | --- | --- | --- | --- | --- | --- |
| corals | 0 | -32% | 0 |  | 0 | Aragonite | High |
| molluscs | -34% | -40% | -17% | -25% |  | Aragonite | High |
| echinoderms | 0 | 0 | -10% | -11% |  | HMC | Medium |
| coccolithophores |  | -23% | 0 |  | 0 | Calcite | Low |
| crustaceans | 0 | 0 | 0 | 0 |  | Calcite | Low |
| calcified algae |  |  | 0 |  | -28% | HMC | Medium |

### 5.1 Protection and adaptation to ocean acidification

Marine calcifiers such as crustose coralline red algae, sea urchins and some molluscan taxa possess skeletons/ shells covered by external organic layers that provide protection from ambient seawater, making them less vulnerable to OA than other taxa (Ries et al., 2009). For instance, crustaceans generally have exoskeletons which are enclosed within epicuticles (Ries et al., 2009). The pteropod *L. helicina antarctica* also has an outer organic layer (periostracum) that can protect the outer shell surface from dissolution (Bednaršek et al., 2012, 2014), as do other molluscs. However, in benthic bivalves such as *L. elliptica*, this periostracum can be easily damaged, exposing the underlying shell exposed to surrounding seawater.

Plasticity in, and control over, biomineralization is a likely mechanism by which calcifiers can adapt to OA. For example, *L. elliptica* showed upregulation in expression of chitin synthase, a gene involved in the shell formation process in response to reduced pH conditions (Cummings et al., 2011). Potentially higher energetic costs of this upregulation were indicated by elevated respiration rates which, when modelled at the population level, predicted significant declines in abundance (Guy et al., 2014). A study of *Mytilus* spp. from temperate to high Arctic latitudes revealed mussels producing thinner shells with a higher organic content in the polar region (Telesca et al., 2019). The periostracum was thicker, and the shells contained more calcite in their prismatic layer than aragonite, which potentially provide greater protection against dissolution (Telesca et al., 2019). The brachiopod *Liothyrella uva* also has the ability to form thicker shells as a compensatory mechanism to dissolution under acidified conditions (Cross et al., 2019). However, the energetic cost of calcification in cold waters may also lead to other compensatory responses. For example, the Antarctic king crab *P. birsteini* invest more resources in building robust, predatory chelae than in protective carapaces, suggesting a greater investment in predation than in defence as its predation pressure is limited in Antarctic waters (Steffel et al., 2019).

### 5.2 Capacity to regulate calcification

Biocalcification (a biologically facilitated process) is either extracellular through deposition of CaCO_3_ on the exterior of an organism; or intracellular and controlled from within the organism (Weiner and Dove, 2003). Extracellular mineralization of CaCO_3_ requires the Ω to be high enough for calcification; and for the organism to control the hydrogen ion concentration in the surrounding area to prevent it bonding with CO_3_^2−^, which reduces the availability of carbonate for shell building. Through the active pumping of CO_3_^2−^ out of a cell, an organism increases the amount of free CO_3_^2−^ for calcification. Alternatively, CO_3_^2−^ can be pumped into a vesicle within a cell and then the vesicle containing the CaCO_3_ is secreted to the outside the cell (Weiner and Dove, 2003). Molluscs and corals generally utilize extracellular calcification secreting their shells and skeletons, respectively, from a calcifying fluid (Ries, 2011).

Intracellular mineralization of CaCO_3_ is achieved within an organism and retained internally to form a skeleton or internal structure, or is secreted to the outside of the organism where it is protected by the covering cell membrane (Weiner and Dove, 2003). Examples of organisms that utilize intracellular calcification include echinoderms and coccolithophores. In particular, coccolithophores control calcification through production of calcified scales (coccoliths) in an intracellular compartment and secrete them to the cell surface (Marsh, 2003). The capacity to regulate calcification via regulation of the pH of intracellular calcifying fluids used to secrete CaCO_3_ may provide some resistance to OA effects and resilience to recover from periods of low pH such as in winter. Some species, such as some corals, can regulate the pH of their calcifying fluid and thus they could buffer OA effects (Ries et al., 2009). Molluscs are generally not capable of regulating the pH and carbonate chemistry of their calcifying fluid (Ries, 2012; Bednaršek et al., 2014). There have been no studies of crustose coralline algae and OA (either distributions, or potential responses) in SO regions to date but the Arctic *Phymatolithon* spp. exhibits strong biological control over their surface chemistry, and efficient carbon concentrating mechanisms, that may prevent net dissolution by elevating pH and CO_3_^2−^ at their surface (Hofmann et al., 2018).

## 6. Predictions about how ocean acidification will affect different pelagic and benthic taxa

It is anticipated that marine calcifiers living in SO regions will be exposed to undersaturated seawater conditions with respect to aragonite by 2050 (Orr et al., 2005; McNeil and Matear, 2008) and with respect to calcite by 2095 (McNeil and Matear, 2008). The situation of some regions south of the Polar Front may be aggravated during austral winters (Orr et al., 2005), when wintertime aragonite undersaturation may occur as early as 2030 in the latitudinal band between 65 and 70°S (McNeil and Matear, 2008; McNeil et al., 2010; Mattsdotter Björk et al., 2014), and during the seasonal decline in primary production, when there is less biological drawdown of CO_2_.

As the saturation horizon shoals, deep water calcifying species that currently live above the saturation horizon may be amongst the earliest to be affected by OA (Turley et al., 2007). Many benthic organisms could also potentially lose their habitat, particularly those on the narrow shallow water shelves of the eastern Weddell Sea and deeper shelves of the Amundsen, Bellingshausen and western Weddell Seas with high contents of CaCO_3_ sediments. As organic matter alters carbonate chemistry of sediments, species inhabiting regions where the flux of organic matter to the seafloor is predicted to increase (e.g. new ice-free areas) could also be at risk. Although taxa characterized by circumpolar distributions and/or broad bathymetric ranges (e.g. *Laternulla elliptica*, *Glabraster antarctica* and *Sterechinus neumayeri*; Fig. 4) could find temporary refuge in shallower waters, and/or in some SO areas where undersaturation may occur later (e.g. far from latitudinal band between 65 and 70°S), it is unclear as distribution shifts will be likely influenced by other environmental factors such as warming.

### 6.1 Mobile organisms

Among molluscs, pteropods may be particularly vulnerable to the projected emergence of a shallow saturation horizon since they generally live in the upper 300 m (Hunt et al., 2008). A recent review and meta-analysis indicated that pteropods are at risk in the Ω_Ar_ range from 1.5–0.9, and that OA effects on survival and shell dissolution are exacerbated with warming (Bednaršek et al., 2019). Experimental studies on the effects of OA on calcifying organisms have focused on a few particular cold-water species. Our meta-analysis shows significant effects of OA on growth (Hoshijima et al., 2017) and shell state (Bednaršek et al., 2014) in the pteropod *Limacina helicina* (Fig. S1). This species was already considered an OA sentinel due to its thin aragonitic shell, which is susceptible to dissolution (Fig. 5) (Bednaršek et al., 2012, 2014). *L. helicina* may also not also have the genetic plasticity required to adapt to OA over the timescales in which the SO is changing (Johnson and Hofmann, 2017). The larvae of the bivalve *Laternula elliptica*, which have predominantly aragonite shells, also seem to be sensitive to undersaturated conditions (shell dissolution and slower development) (Fig. S1) (Bylenga et al., 2015, 2017). Given its circumpolar distribution (Fig. 4) and broad depth range (<10 m – ~700 m) (Waller et al., 2017), individuals from shallower waters could escape from initial predicted occurrences of aragonite undersaturation events.

**Fig. 5.**
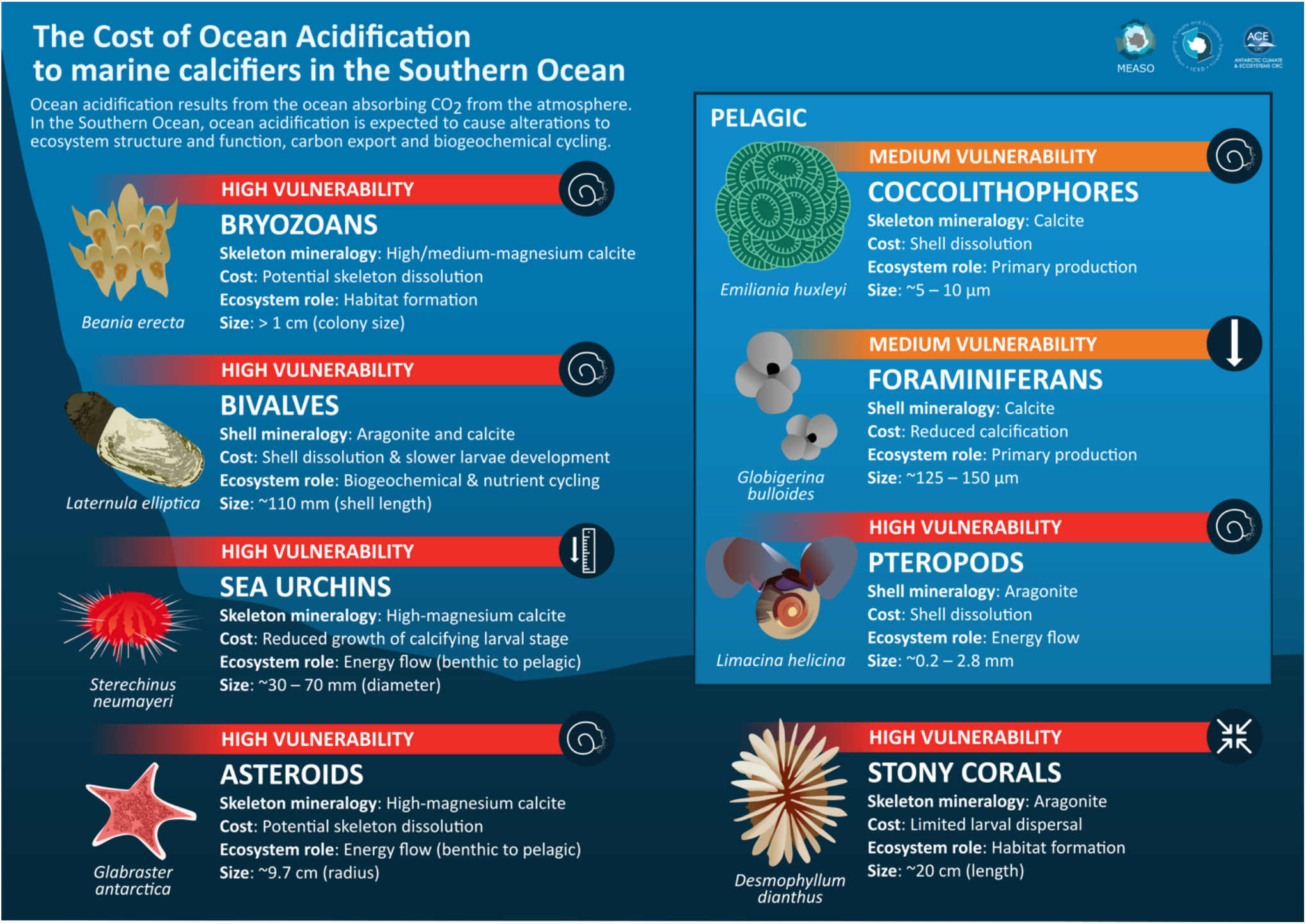
Infographic of predictions about how ocean acidification will affect particular groups of marine calcifiers from Antarctica. Information on different mineral composition, functional and life history traits is provided for each group.

Information compiled here also reveals that sea stars and brittle stars may be highly vulnerable to OA due to their HMC skeletons, and the apparent lack of strategies to overcome reduced pH/carbonate conditions (Table 2). The negative impact of OA on echinoderms, especially on some sea star species which are common key predators in Antarctic ecosystems, could alter the structure and biodiversity of these ecosystems, with cascading impacts on multiple trophic levels. One example is the sea star *Glabraster antarctica*, which has higher mean Mg values (>10 wt% MgCO_3_) than other SO species. Gonzalez-Bernat et al. (2013) showed the growth of the larvae of the seastar *Odontaster validus* is significantly affected when exposed to low pH levels. Previous studies indicate differential sensitivities depending on the biological responses in the common Antarctic sea urchin *Sterechinus neumayeri*. While its fertilization success and early development were not affected by near-future changes, the former declined significantly when only exposed to low pH levels (Ericson et al., 2010; Ho et al., 2013). In addition, other studies have revealed significant interactive effects of projected changes in seawater temperature and pH on the percentage of fertilization (Ericson et al., 2012) and on the cleavage success (Foo et al., 2016) although sperm concentrations, population or experimental conditions have to be considered (Sewell et al., 2014). The growth of *S. neumayeri*’s calcifying larval stage was also found to be negatively correlated with reduced pH and Ω (Byrne et al., 2013) although its calcification was not reduced in a previous study (Clark et al., 2009). This sensitivity of larvae could cause a key bottleneck in their life-cycle (Dupont et al., 2010). Both species are circumpolar and exhibit a high range of eurybathy, ranging from the shallow subtidal to 2,930 m (*O. validus*) and to 500 m depth (*S. neymayeri*) (Pierrat et al., 2012; Moore et al., 2018)). Like other SO echinoderm taxa, the distribution of these common species may become limited to shallow waters above the ASH, likely being then affected by warming.

### 6.2 Sessile organisms

Corals, bryozoans and spirorbid polychaetes are ecosystem engineers, creating habitat complexity and being essential refugia for many associated macro- and micro-invertebrates. Given their biological traits (adult sessile organisms with limited larval dispersal) (Thatje, 2012), their distribution is less likely to shift to more favourable habitat. Dissolution effects could thus lead to loss of habitat complexity with serious consequences to the Antarctic ecosystems. Spirorbid polychaetes may be highly vulnerable to OA due to their HMC and/or aragonitic skeletons (Cigliano et al., 2010; Smith et al., 2013b; Peck et al., 2015) and a lack a protective external organic sheet equivalent to molluscan periostracum (Tanur et al., 2010).

Among corals, the scleractinid coral species *Desmophyllum dianthus*, *Solenosmilia sp.* and *Madrepora* sp., with aragonitic skeletons, will also probably be impacted due to their mineralogy-related sensitivity and their biological traits. *D. dianthus* has a common depth range between 200– 2500 m (Miller et al., 2011; Fillinger and Richter, 2013), and exhibits reduced respiration rates under predicted combination of rising temperatures and OA scenarios (Gori et al., 2016). Thus it is likely this species may be severely impacted by these two key CO_2_-related stressors. Scleractinid corals comprise dominant components of VMEs on the Antarctic shelf and the effects of OA at the species level could thus potentially lead to wider ecosystem level effects. However, there is a need to better understand the long-term implications and mechanisms of OA. For example, the congeneric deep-sea species *Solenosmilia variabilis* from New Zealand (3.5°C) maintained for 12 months under reduced pH conditions (pH = 7.65, Ω_Ar_ = 0.69 ± 0.01) and only showed colony tissue loss (Gammon et al., 2018). The authors hypothesized that this response could indicate a reallocation of energy, with physiological processes (e.g. growth and respiration) being maintained at the expense of coenenchyme production (the common tissue that surrounds and links the polyps in octocorals) (Gammon et al., 2018).

Another sessile species, the common bryozoan *Beania erecta*, could also be susceptible to OA conditions as some individuals were found to secrete HMC skeletons, even though the range was highly variable (5.3–13.6 wt% MgCO_3_ in calcite) (Figuerola et al., 2019). *B. erecta*, like other Antarctic bryozoan species, can form dense mats that, in some cases, cover large areas of substratum. Potential decreases of some species could indirectly negatively impact other organisms which use their colonies as substrate, food and shelter (Figuerola et al., 2019).

## 7 Future directions

The SO may serve as a sentinel for evaluating the impacts of OA on marine calcifiers and aid in making predictions of the future impacts at lower latitudes, where OA is expected to occur later. Some SO regions are also warming faster than other global oceans, along with other stressors such as nutrient inputs. Key marine calcifiers may not be able to cope with these multiple simultaneous projected changes, which could ultimately affect food webs and have cascading impacts from low to higher trophic levels. Combined temperature and pH studies of several Antarctic taxa indicate that temperature may have a stronger effect on survival and growth, but that effects of OA may be severe (e.g. increased rates of abnormal development, sublethal effects). Such studies of multiple stressors are essential to explore the capacity of different species to adapt to future environmental changes. However, our understanding on the potential impact of OA on Antarctic marine calcifiers, including keystone organisms, is still constrained by the limited mineralogical and biological data available for most species. Furthermore, broader understanding is also constrained by the limited geographical coverage of studies, lack of reliable long-term monitoring, and the overall lack of multidisciplinary studies over longer times scales integrating interactions between stressors and an array of species with different mineral composition, functional and life history traits. In particular, other environmental and biological factors, which may influence mineral susceptibility (such as organic content, protective periostracum, physiological constraints in cold waters), need further evaluation. Understanding of ecosystem effects can be improved through the use of *in-situ* OA studies over long times scales, such as by using free-ocean CO_2_ enrichment (FOCE) systems (Stark et al., 2018). Being semi-open systems, these not only allow manipulation of pH, but also incorporate natural variation in physicochemical variables, such as temperature and salinity, thus creating realistic near-future OA conditions. Recently, FOCE technology has successfully been implemented in Antarctica (Stark et al., 2018) and other *in situ* OA experiments have been conducted (Barr et al., 2017; Cummings et al., 2019), opening new possibilities for OA research in polar regions. Finally, special attention should be paid to the use of standardized protocols for experimental approaches, the manipulation and reporting carbonate chemistry conditions (Riebesell et al., 2010), in order to usefully in order to compare results from different studies and make predictions about how OA may impact different groups.

Recent studies have examined relationships between variations in the mineralogical composition of Antarctic and Subantarctic marine calcifiers and decreasing seawater pH and greater solubility of HMC and aragonite. However, the expected decrease of skeletal Mg-calcite along a depth gradient in four Antarctic bryozoan species was not apparent (Figuerola et al., 2015), and nor was an expected increase of aragonitic forms towards shallower waters in sub-Antarctic stylasterid populations (Bax and Robinson, unpublished data). These mineralogy findings do not consider other factors influencing the species distributions, such as gene flow across depth gradients which may work against selective pressures influencing mineralogy (Miller et al., 2011). Our meta-analysis highlights the detrimental impact of OA on dispersal stages which could have significant long-term consequences for isolated populations (Miller et al., 2011) or eventually lead to depth related differences in mineralogy. A further step will be to consider a broader array of species and a wider depth range to test the effect of the lower CaCO_3_ Ω on distributions. There is a need for information on mineralogy of all major groups of calcifying invertebrates over a wider depth range to evaluate patterns related to environmental (e.g. local CSH and ASH) and biological factors (e.g. food availability), with a focus on identifying taxa or communities that may be potentially vulnerable to near-future OA, and that may make suitable indicators (sentinels) to monitor effects of OA in the SO. Building interdisciplinary collaborations via projects such as the Census of Antarctic Marine Life (CAML) is essential (Gutt et al., 2018) to successfully perform OA research on and in remote and at risk regions like Antarctica.

## Acknowledgments

We thank to Stacey McCormack for developing Figures 1 and 5. JS publishes with the permission of the CEO, Geoscience Australia.

## Author Contributions

BF conceived the initial idea for the review, organized the sections and the database and made tables for the mineralogical data, wrote the first draft of the manuscript and edited the final version. BF and AC provided the framework for the document and BF, AH, NB and AC participated in the design advice for the figures. HG reviewed and analysed the data available to produce species distribution maps. AH and BF screened papers and AH performed the meta-analysis and its figures. AH and BF interpreted the results. Authors reviewed the first draft with substantial contributions by section as follows: AH (2, 3 and 5: meta-analysis), NB (4.1: corals; 7), VC (4.2: algae and echinoderms; 5.1), RD (4.2: sponges), and JS (1; 1.1.2: carbonate sediments) and JSS (1; 5.2). All authors provided comments and suggestions and approved the final manuscript.

## References

Allcock, A. L., and Strugnell, J. M. (2012). Southern Ocean diversity: new paradigms from molecular ecology. Trends Ecol. Evol. 27, 520–528. doi:10.1016/j.tree.2012.05.009.

Alongi, G., Cormaci, M., and Furnari, G. (2002). The Corallinaceae (Rhodophyta) from the Ross Sea (Antarctica): a taxonomic revision rejects all records except *Phymatolithon foecundum*. Phycologia 41, 140–146. doi:10.2216/i0031-8884-41-2-140.1.

Anand, P., and Elderfield, H. (2005). Variability of Mg/Ca and Sr/Ca between and within the planktonic foraminifers *Globigerina bulloides* and *Globorotalia truncatulinoides*. Geochem. Geophys. Geosystems 6. doi:10.1029/2004GC000811.

Andersson, A. J., and Gledhill, D. (2013). Ocean Acidification and Coral Reefs: Effects on Breakdown, Dissolution, and Net Ecosystem Calcification. Annu. Rev. Mar. Sci. 5, 321–348. doi:10.1146/annurev-marine-121211-172241.

Andersson, A. J., and Mackenzie, F. T. (2011). Technical comment on Kroeker et al. (2010) Meta-analysis reveals negative yet variable effects of ocean acidification on marine organisms. Ecology Letters, 13, 1419–1434. Ecol. Lett. 14, E1–E2. doi:10.1111/j.1461-0248.2011.01646.x.

Andersson, A. J., Mackenzie, F. T., and Bates, N. R. (2008). Life on the margin: Implications of ocean acidification on Mg-calcite, high latitude and cold-water marine calcifiers. Mar. Ecol. Prog. Ser. 373, 265–273. doi:10.3354/meps07639.

Baird, H. P., Miller, K. J., and Stark, J. S. (2011). Evidence of hidden biodiversity, ongoing speciation and diverse patterns of genetic structure in giant Antarctic amphipods. Mol. Ecol. 20, 3439–3454. doi:10.1111/j.1365-294X.2011.05173.x.

Barker, S., and Elderfield, H. (2002). Foraminiferal Calcification Response to Glacial-Interglacial Changes in Atmospheric CO2. Science 297, 833–836. doi:10.1126/science.1072815.

Barnes, D. K. A., and Kuklinski, P. (2010). Bryozoans of the Weddell Sea continental shelf, slope and abyss: did marine life colonize the Antarctic shelf from deep water, outlying islands or in situ refugia following glaciations? J. Biogeogr. 37, 1648–1656. doi:10.1111/j.1365-2699.2010.02320.x.

Barnes, D. K. A., and Peck, L. S. (1996). Epibiota and attachment substrata of deep-water brachiopods from Antarctica and New Zealand. Philos. Trans. R. Soc. Lond. B. Biol. Sci. 351, 677–687. doi:10.1098/rstb.1996.0064.

Barnes, D. K. A., and Peck, L. S. (2008). Vulnerability of Antarctic shelf biodiversity to predicted regional warming. Clim. Res. 37, 149–163. doi:10.3354/cr00760.

Barr, N. G., Lohrer, A. M., and Cummings, V. J. (2017). An *in situ* incubation method for measuring the productivity and responses of under-ice algae to ocean acidification and warming in polar marine habitats. Limnol. Oceanogr. Methods 15, 264–275. doi:10.1002/lom3.10154.

Bax, N. (2015). Deep-Sea Stylasterid Corals in the Antarctic, SubAntarctic and Patagonian Benthos: Biogeography, Phylogenetics, Connectivity and Conservation. Available at: https://eprints.utas.edu.au/22768/2/whole_Bax_thesis_ex_pub_mat.pdf.

Bax, N. N., and Cairns, S. D. (2014). “Stylasteridae (Cnidaria; Hydrozoa),” in DeBroyer C et al. Biogeographic Atlas of the Southern Ocean. Cambridge: SCAR 107-112 (SCAR). Available at: http://repository.si.edu/xmlui/handle/10088/22593 [Accessed June 11, 2020].

Beaman, R. J., and Harris, P. T. (2005). Bioregionalisation of the George V Shelf, East Antarctica. Cont. Shelf Res. 25, 1657–1691. doi:10.1016/j.csr.2005.04.013.

Beaufort, L., Probert, I., de Garidel-Thoron, T., Bendif, E. M., Ruiz-Pino, D., Metzl, N., et al. (2011). Sensitivity of coccolithophores to carbonate chemistry and ocean acidification. Nature 476, 80–83. doi:10.1038/nature10295.

Bednaršek, N., Feely, R. A., Howes, E. L., Hunt, B. P. V., Kessouri, F., León, P., et al. (2019). Systematic Review and Meta-Analysis Toward Synthesis of Thresholds of Ocean Acidification Impacts on Calcifying Pteropods and Interactions With Warming. Front. Mar. Sci. 6, 227. doi:10.3389/fmars.2019.00227.

Bednaršek, N., Tarling, G. A., Bakker, D. C. E., Fielding, S., and Feely, R. A. (2014). Dissolution Dominating Calcification Process in Polar Pteropods Close to the Point of Aragonite Undersaturation. PLOS ONE 9, e109183. doi:10.1371/journal.pone.0109183.

Bednaršek, N., Tarling, G. A., Bakker, D. C. E., Fielding, S., Jones, E. M., Venables, H. J., et al. (2012). Extensive dissolution of live pteropods in the Southern Ocean. Nat. Geosci. 5, 881–885. doi:10.1038/ngeo1635.

Blackmon, P. D., and Todd, R. (1959). Mineralogy of Some Foraminifera as Related to Their Classification and Ecology. J. Paleontol. 33, 1–15.

Borisenko, Y., and Gontar, V. I. (1991). Biogeochemistry of skeletons of coldwater Bryozoa. Biol Morya 1, 80–90.

Brandt, A., Gooday, A. J., Brandão, S. N., Brix, S., Brökeland, W., Cedhagen, T., et al. (2007). First insights into the biodiversity and biogeography of the Southern Ocean deep sea. Nature 447, 307–11. doi:10.1038/nature05827.

Bylenga, C. H., Cummings, V. J., and Ryan, K. G. (2015). Fertilisation and larval development in an Antarctic bivalve, *Laternula elliptica*, under reduced pH and elevated temperatures. Mar. Ecol. Prog. Ser. 536, 187–201. doi:10.3354/meps11436.

Bylenga, C. H., Cummings, V. J., and Ryan, K. G. (2017). High resolution microscopy reveals significant impacts of ocean acidification and warming on larval shell development in *Laternula elliptica*. PLOS ONE 12, e0175706. doi:10.1371/journal.pone.0175706.

Byrne, M., Ho, M. A., Koleits, L., Price, C., King, C. K., Virtue, P., et al. (2013). Vulnerability of the calcifying larval stage of the Antarctic sea urchin *Sterechinus neumayeri* to near-future ocean acidification and warming. Glob. Change Biol. 19, 2264–2275. doi:10.1111/gcb.12190.

Cairns, S. D. (1982). Antarctic and Subantarctic Scleractinia. Antarct. Res. Ser. 34, 1–74. doi:10.1029/AR034p0001.

Cairns, S. D. (1983). A Generic Revision of the Stylasterina (Coelenterata: Hydrozoa). Part 1. Description of the Genera. Bull. Mar. Sci. 33, 427–508.

Cairns, S. D., and Macintyre, I. G. (1992). Phylogenetic Implications of Calcium Carbonate Mineralogy in the Stylasteridae (Cnidaria: Hydrozoa). PALAIOS 7, 96–107. doi:10.2307/3514799.

Cairns, S. D., and Polonio, V. (2013). New records of deep-water Scleractinia off Argentina and the Falkland Islands. Zootaxa 3691, 58–86. doi:10.11646/zootaxa.3691.1.2.

Chave, K. E. (1954). Aspects of the Biogeochemistry of Magnesium 1. Calcareous Marine Organisms. J. Geol. 62, 266–283. doi:10.1086/626162.

Cigliano, M., Gambi, M. C., Rodolfo-Metalpa, R., Patti, F. P., and Hall-Spencer, J. M. (2010). Effects of ocean acidification on invertebrate settlement at volcanic CO_2_ vents. Mar. Biol. 157, 2489–2502. doi:10.1007/s00227-010-1513-6.

Clark, D., Lamare, M., and Barker, M. (2009). Response of sea urchin pluteus larvae (Echinodermata: Echinoidea) to reduced seawater pH: a comparison among a tropical, temperate, and a polar species. Mar. Biol. 156, 1125–1137. doi:10.1007/s00227-009-1155-8.

Collard, M., Ridder, C. D., David, B., Dehairs, F., and Dubois, P. (2015). Could the acid–base status of Antarctic sea urchins indicate a better-than-expected resilience to near-future ocean acidification? Glob. Change Biol. 21, 605–617. doi:10.1111/gcb.12735.

Conrad, C. J., and Lovenduski, N. S. (2015). Climate-Driven Variability in the Southern Ocean Carbonate System. J. Clim. 28, 5335–5350. doi:10.1175/JCLI-D-14-00481.1.

Cormaci, M., Furnari, G., and Scammacca, B. (2000). “The Macrophytobenthos of Terra Nova Bay,” in Ross Sea Ecology: Italiantartide Expeditions (1987–1995), eds. F. M. Faranda, L. Guglielmo, and A. Ianora (Berlin, Heidelberg: Springer), 493–502. doi:10.1007/978-3-642-59607-0_35.

Cross, E. L., Harper, E. M., and Peck, L. S. (2019). Thicker Shells Compensate Extensive Dissolution in Brachiopods under Future Ocean Acidification. Environ. Sci. Technol. 53, 5016–5026. doi:10.1021/acs.est.9b00714.

Cross, E. L., Peck, L. S., and Harper, L. (2014). Ocean acidification does not impact shell growth or repair of the Antarctic brachiopod *Liothyrella uva* (Broderip, 1833). doi:https://www.repository.cam.ac.uk/handle/1810/247599.

Cummings, V., Hewitt, J., Rooyen, A. V., Currie, K., Beard, S., Thrush, S., et al. (2011). Ocean Acidification at High Latitudes: Potential Effects on Functioning of the Antarctic Bivalve *Laternula elliptica*. PLOS ONE 6, e16069. doi:10.1371/journal.pone.0016069.

Cummings, V. J., Barr, N. G., Budd, R. G., Marriott, P. M., Safi, K. A., and Lohrer, A. M. (2019). In situ response of Antarctic under-ice primary producers to experimentally altered pH. Sci. Rep. 9, 6069. doi:10.1038/s41598-019-42329-0.

Cyronak, T., Schulz, K. G., and Jokiel, P. L. (2016). The Omega myth: what really drives lower calcification rates in an acidifying ocean. ICES J. Mar. Sci. 73, 558–562. doi:10.1093/icesjms/fsv075.

Davis, C. V., Rivest, E. B., Hill, T. M., Gaylord, B., Russell, A. D., and Sanford, E. (2017). Ocean acidification compromises a planktic calcifier with implications for global carbon cycling. Sci. Rep. 7, 1–8. doi:10.1038/s41598-017-01530-9.

Dayton, P. K., Robilliard, G. A., Paine, R. T., and Dayton, L. B. (1974). Biological Accommodation in the Benthic Community at McMurdo Sound, Antarctica. Ecol. Monogr. 44, 105–128.

Donahue, K., Klaas, C., Dillingham, P. W., and Hoffmann, L. J. (2019). Combined effects of ocean acidification and increased light intensity on natural phytoplankton communities from two Southern Ocean water masses. J. Plankton Res. 41, 30–45. doi:10.1093/plankt/fby048.

Downey, R. V., Griffiths, H. J., Linse, K., and Janussen, D. (2012). Diversity and Distribution Patterns in High Southern Latitude Sponges. PLOS ONE 7, e41672. doi:10.1371/journal.pone.0041672.

Dupont, S., Dorey, N., and Thorndyke, M. (2010). What meta-analysis can tell us about vulnerability of marine biodiversity to ocean acidification? Estuar. Coast. Shelf Sci. 89, 182–185. doi:10.1016/j.ecss.2010.06.013.

Duquette, A., Halanych, K. M., Angus, R. A., and McClintock, J. B. (2018). Inter and intraspecific comparisons of the skeletal Mg/Ca ratios of high latitude Antarctic echinoderms. Antarct. Sci. 30, 160–169. doi:10.1017/S0954102017000566.

Ericson, J. A., Ho, M. A., Miskelly, A., King, C. K., Virtue, P., Tilbrook, B., et al. (2012). Combined effects of two ocean change stressors, warming and acidification, on fertilization and early development of the Antarctic echinoid *Sterechinus neumayeri*. Polar Biol. 35, 1027–1034. doi:10.1007/s00300-011-1150-7.

Ericson, J. A., Lamare, M. D., Morley, S. A., and Barker, M. F. (2010). The response of two ecologically important Antarctic invertebrates (*Sterechinus neumayeri* and *Parborlasia corrugatus*) to reduced seawater pH: effects on fertilisation and embryonic development. Mar. Biol. 157, 2689–2702. doi:10.1007/s00227-010-1529-y.

Fabry, V. J. (2008). Marine Calcifiers in a High-CO₂ Ocean. Science 320, 1020–1022. doi:10.1126/science.1157130.

Fabry, V., McClintock, J., Mathis, J., and Grebmeier, J. (2009). Ocean Acidification at High Latitudes: The Bellwether. Oceanography 22, 160–171. doi:10.5670/oceanog.2009.105.

Feely, R. A., Sabine, C. L., Lee, K., Berelson, W., Kleypas, J., Fabry, V. J., et al. (2004). Impact of anthropogenic CO_2_ on the CaCO_3_ system in the oceans. Science 305, 362–366. doi:10.1126/science.1097329.

Figuerola, B., Ballesteros, M., Monleón-Getino, T., and Avila, C. (2012). Spatial patterns and diversity of bryozoan communities from the Southern Ocean: South Shetland Islands, Bouvet Island and Eastern Weddell Sea. Syst. Biodivers. 10, 109–123. doi:10.1080/14772000.2012.668972.

Figuerola, B., Gore, D. B., Johnstone, G., and Stark, J. S. (2019). Spatio-temporal variation of skeletal Mg-calcite in Antarctic marine calcifiers. PLOS ONE 14, e0210231. doi:10.1371/journal.pone.0210231.

Figuerola, B., Kuklinski, P., Carmona, F., and Taylor, P. D. (2017). Evaluating potential factors influencing branch diameter and skeletal Mg-calcite using an Antarctic cyclostome bryozoan species. Hydrobiologia 799. doi:10.1007/s10750-017-3213-4.

Figuerola, B., Kuklinski, P., and Taylor, P. D. (2015). Depth patterns in Antarctic bryozoan skeletal Mg-calcite: Can they provide an analogue for future environmental changes? Mar. Ecol. Prog. Ser. 540, 109–120. doi:10.3354/meps11515.

Fillinger, L., and Richter, C. (2013). Vertical and horizontal distribution of *Desmophyllum dianthus* in Comau Fjord, Chile: a cold-water coral thriving at low pH. PeerJ 1, e194. doi:https://doi.org/10.7717/peerj.194.

Foo, S. A., Sparks, K. M., Uthicke, S., Karelitz, S., Barker, M., Byrne, M., et al. (2016). Contributions of genetic and environmental variance in early development of the Antarctic sea urchin *Sterechinus neumayeri* in response to increased ocean temperature and acidification. Mar. Biol. 163, 130. doi:10.1007/s00227-016-2903-1.

Gammon, M. J., Tracey, D. M., Marriott, P. M., Cummings, V. J., and Davy, S. K. (2018). The physiological response of the deep-sea coral *Solenosmilia variabilis* to ocean acidification. PeerJ 6, e5236. doi:10.7717/peerj.5236.

Gonzalez-Bernat, M. J., Lamare, M., and Barker, M. (2013). Effects of reduced seawater pH on fertilisation, embryogenesis and larval development in the Antarctic seastar *Odontaster validus*. Polar Biol. 36, 235–247. doi:10.1007/s00300-012-1255-7.

Gori, A., Ferrier-Pagès, C., Hennige, S. J., Murray, F., Rottier, C., Wicks, L. C., et al. (2016). Physiological response of the cold-water coral *Desmophyllum dianthus* to thermal stress and ocean acidification. PeerJ 4, e1606. doi:10.7717/peerj.1606.

Griffiths, H. J., Whittle, R. J., Roberts, S. J., Belchier, M., and Linse, K. (2013). Antarctic Crabs: Invasion or Endurance? PLOS ONE 8, e66981. doi:10.1371/journal.pone.0066981.

Guinotte, J. M., Orr, J., Cairns, S., Freiwald, A., Morgan, L., and George, R. (2006). Will human-induced changes in seawater chemistry alter the distribution of deep-sea scleractinian corals? Front. Ecol. Environ. 4, 141–146. doi:10.1890/1540-9295(2006)004[0141:WHCISC]2.0.CO;2.

Gutt, J., Isla, E., Bertler, A. N., Bodeker, G. E., Bracegirdle, T. J., Cavanagh, R. D., et al. (2018). Cross-disciplinarity in the advance of Antarctic ecosystem research. Mar. Genomics 37, 1–17. doi:10.1016/j.margen.2017.09.006.

Guy, C. I., Cummings, V. J., Lohrer, A. M., Gamito, S., and Thrush, S. F. (2014). Population trajectories for the Antarctic bivalve *Laternula elliptica*: identifying demographic bottlenecks in differing environmental futures. Polar Biol. 37, 541–553. doi:10.1007/s00300-014-1456-3.

Haese, R. R., Smith, J., Weber, R., and Trafford, J. (2014). High-Magnesium Calcite Dissolution in Tropical Continental Shelf Sediments Controlled by Ocean Acidification. Environ. Sci. Technol. 48, 8522–8528.

Hancock, A. M., King, C. K., Stark, J. S., McMinn, A., and Davidson, A. T. (2020). Effects of ocean acidification on Antarctic marine organisms: A meta-analysis. Ecol. Evol. 00, 1–20. doi:10.1002/ece3.6205.

Hauck, J., Gerdes, D., Hillenbrand, C.-D., Hoppema, M., Kuhn, G., Nehrke, G., et al. (2012). Distribution and mineralogy of carbonate sediments on Antarctic shelves. J. Mar. Syst. 90, 77–87. doi:10.1016/j.jmarsys.2011.09.005.

Hauri, C., Friedrich, T., and Timmermann, A. (2016). Abrupt onset and prolongation of aragonite undersaturation events in the Southern Ocean. Nat. Clim. Change 6, 172–176. doi:10.1038/nclimate2844.

Hedges, L., and Olkin, I. (1985). Statistical Methods for Meta-Analysis. New York, NY: Academic Press doi:10.1016/C2009-0-03396-0.

Hermans, J., Borremans, C., Willenz, P., André, L., and Dubois, P. (2010). Temperature, salinity and growth rate dependences of Mg/Ca and Sr/Ca ratios of the skeleton of the sea urchin *Paracentrotus lividus* (Lamarck): An experimental approach. Mar. Biol. 157, 1293–1300. doi:10.1007/s00227-010-1409-5.

Ho, M. A., Price, C., King, C. K., Virtue, P., and Byrne, M. (2013). Effects of ocean warming and acidification on fertilization in the Antarctic echinoid *Sterechinus neumayeri* across a range of sperm concentrations. Mar. Environ. Res. 90, 136–141. doi:10.1016/j.marenvres.2013.07.007.

Hofmann, L. C., Schoenrock, K., and de Beer, D. (2018). Arctic Coralline Algae Elevate Surface pH and Carbonate in the Dark. Front. Plant Sci. 9. doi:10.3389/fpls.2018.01416.

Hoshijima, U., Wong, J. M., and Hofmann, G. E. (2017). Additive effects of pCO2 and temperature on respiration rates of the Antarctic pteropod *Limacina helicina antarctica*. Conserv. Physiol. 5, cox064. doi:10.1093/conphys/cox064.

Hunt, B. P. V., Pakhomov, E. A., Hosie, G. W., Siegel, V., Ward, P., and Bernard, K. (2008). Pteropods in Southern Ocean ecosystems. Prog. Oceanogr. 78, 193–221. doi:10.1016/j.pocean.2008.06.001.

IPCC (2014). Synthesis report in climate change 2014: contribution of working groups I, II and III to the fifth assessment report of the Intergovernmental Panel on Climate Change. Geneva, Switzerland: IPCC.

IPCC (2019). IPCC special report on the ocean and cryosphere in a changing climate. Geneva, Switzerland Available at: http://www.unenvironment.org/resources/report/ipcc-special-report-ocean-and-cryosphere-changing-climate.

Irving, A. D., Connell, S. D., Johnston, E. L., Pile, A. J., and Gillanders, B. M. (2005). The response of encrusting coralline algae to canopy loss: an independent test of predictions on an Antarctic coast. Mar. Biol. 147, 1075–1083. doi:10.1007/s00227-005-0007-4.

Jantzen, C., Häussermann, V., Försterra, G., Laudien, J., Ardelan, M., Maier, S., et al. (2013). Occurrence of a cold-water coral along natural pH gradients (Patagonia, Chile). Mar. Biol. 160, 2597–2607. doi:10.1007/s00227-013-2254-0.

Janussen, D., and Downey, R. (2014). “Porifera,” in DeBroyer C et al. Biogeographic Atlas of the Southern Ocean. Cambridge: SCAR 107-112 (Cambridge), 94–102.

Jiang, L.-Q., Feely, R. A., Carter, B. R., Greeley, D. J., Gledhill, D. K., and Arzayus, K. M. (2015). Climatological distribution of aragonite saturation state in the global oceans. Glob. Biogeochem. Cycles 29, 1656–1673. doi:10.1002/2015GB005198.

Johnson, K. M., and Hofmann, G. E. (2017). Transcriptomic response of the Antarctic pteropod *Limacina helicina antarctica* to ocean acidification. BMC Genomics 18, 812. doi:10.1186/s12864-017-4161-0.

Karelitz, S. E., Uthicke, S., Foo, S. A., Barker, M. F., Byrne, M., Pecorino, D., et al. (2017). Ocean acidification has little effect on developmental thermal windows of echinoderms from Antarctica to the tropics. Glob. Change Biol. 23, 657–672. doi:10.1111/gcb.13452.

Kleypas, J. A., Buddemeier, R. W., Archer, D., Gattuso, J.-P., Langdon, C., and Opdyke, B. N. (1999). Geochemical Consequences of Increased Atmospheric Carbon Dioxide on Coral Reefs. Science 284, 118–120. doi:10.1126/science.284.5411.118.

Kroeker, K. J., Kordas, R. L., Crim, R., Hendriks, I. E., Ramajo, L., Singh, G. S., et al. (2013). Impacts of ocean acidification on marine organisms: quantifying sensitivities and interaction with warming. Glob. Change Biol. 19, 1884–1896. doi:10.1111/gcb.12179.

Kroeker, K. J., Kordas, R. L., Crim, R. N., and Singh, G. G. (2010). Meta-analysis reveals negative yet variable effects of ocean acidification on marine organisms. Ecol. Lett. 13, 1419–1434. doi:10.1111/j.1461-0248.2010.01518.x.

Kroeker, K. J., Kordas, R. L., Crim, R. N., and Singh, G. G. (2011). Response to technical comment on ‘meta-analysis reveals negative yet variable effects of ocean acidification on marine organisms’. *Ecol. Lett.* 14, E1–E2. doi:10.1111/j.1461-0248.2011.01665.x.

Krzeminska, M., Kuklinski, P., Najorka, J., and Iglikowska, A. (2016). Skeletal Mineralogy Patterns of Antarctic Bryozoa. J. Geol. 124, 411–422. doi:10.1086/685507.

Lebrato, M., Andersson, A. J., Ries, J. B., Aronson, R. B., Lamare, M. D., Koeve, W., et al. (2016). Benthic marine calcifiers coexist with CaCO_3_-undersaturated seawater worldwide. Glob. Biogeochem. Cycles 30, 1038–1053. doi:10.1002/2015GB005260.

Lebrato, M., Feely, R. A., Greeley, D., Jones, O. B., Suarez-bosche, N., Lampitt, R. S., et al. (2010). Global contribution of echinoderms to the marine carbon cycle: a re-assessment of the oceanic CaCO_3_ budget and the benthic compartments. Ecol. Monogr. 80, 441–467.

Lohbeck, K. T., Riebesell, U., and Reusch, T. B. H. (2012). Adaptive evolution of a key phytoplankton species to ocean acidification. Nat. Geosci. 5, 346–351. doi:10.1038/ngeo1441.

Lowenstam, H. A. (1954). Environmental Relations of Modification Compositions of Certain Carbonate Secreting Marine Invertebrates. Proc. Natl. Acad. Sci. 40, 39–48. doi:10.1073/pnas.40.1.39.

Loxton, J., Kuklinski, P., Barnes, D. K. A., Najorka, J., Jones, M. S., and Porter, J. S. (2014). Variability of Mg-calcite in Antarctic bryozoan skeletons across spatial scales. Mar. Ecol. Prog. Ser. 507, 169–180. doi:10.3354/meps10826.

Loxton, J., Kuklinski, P., Mair, J. M., Jones, M. S., and Porter, J. S. (2013). Patterns of Magnesium-Calcite Distribution in the Skeleton of Some Polar Bryozoan Species Mineralogy of Polar Bryozoan Skeletons. Bryozoan Stud. 2010 143, 169–185. doi:10.1007/978-3-642-16411-8.

Mackenzie, F. T., Bischoff, W. D., Bishop, F. C., Loijens, M., Schoonmaker, J., and Wollast, R. (1983). “Magnesian calcites: low-temperature occurrence, solubility and solid-solution behavior,” in Carbonates: Mineralogy and Chemistry. Reviews in Mineralogy, ed. R. Reeder (Mineralogical Society of America, Washington, DC), 97–143.

Manno, C., Morata, N., and Bellerby, R. (2012). Effect of ocean acidification and temperature increase on the planktonic foraminifer *Neogloboquadrina pachyderma*(sinistral). Polar Biol. 35, 1311–1319. doi:10.1007/s00300-012-1174-7.

Margolin, A. R., Robinson, L. F., Burke, A., Waller, R. G., Scanlon, K. M., Roberts, M. L., et al. (2014). Temporal and spatial distributions of cold-water corals in the Drake Passage: Insights from the last 35,000 years. Deep Sea Res. Part II Top. Stud. Oceanogr. 99, 237–248. doi:10.1016/j.dsr2.2013.06.008.

Marsh, M. E. (2003). Regulation of CaCO3 formation in coccolithophores. Comp. Biochem. Physiol. B Biochem. Mol. Biol. 136, 743–754. doi:10.1016/s1096-4959(03)00180-5.

Marshall, B. J., Thunell, R. C., Henehan, M. J., Astor, Y., and Wejnert, K. E. (2013). Planktonic foraminiferal area density as a proxy for carbonate ion concentration: A calibration study using the Cariaco Basin ocean time series. Paleoceanography 28, 363–376. doi:http://dx.doi.org/10.1002/palo.20034.

Mattsdotter Björk, M., Fransson, A., Torstensson, A., and Chierici, M. (2014). Ocean acidification state in western Antarctic surface waters: controls and interannual variability. Biogeosciences 11, 57–73. doi:https://doi.org/10.5194/bg-11-57-2014.

McClintock, J. B., Amsler, M. O., Angus, R. A., Challener, R. C., Schram, J. B., Amsler, C. D., et al. (2011). The Mg-Calcite composition of Antarctic echinoderms: important implications for predicting the impacts of ocean acidification. J. Geol. 119, 457–466. doi:10.1086/660890.

McClintock, J. B., Angus, R. A., Mcdonald, M. R., Amsler, C. D., Catledge, S. A., and Vohra, Y. K. (2009). Rapid dissolution of shells of weakly calcified Antarctic benthic macroorganisms indicates high vulnerability to ocean acidification. Antarct. Sci. 21, 449–456. doi:10.1017/S0954102009990198.

McFadden, C. S., Sánchez, J. A., and France, S. C. (2010). Molecular phylogenetic insights into the evolution of Octocorallia: a review. Integr. Comp. Biol. 50, 389–410. doi:10.1093/icb/icq056.

McNeil, B. I., and Matear, R. J. (2008). Southern Ocean acidification: A tipping point at 450-ppm atmospheric CO2. Proc. Natl. Acad. Sci. U. S. A. 105, 18860–18864.

McNeil, B. I., Tagliabue, A., and Sweeney, C. (2010). A multi-decadal delay in the onset of corrosive ‘acidified’ waters in the Ross Sea of Antarctica due to strong air-sea CO_2_ disequilibrium. Geophys. Res. Lett. 37. doi:10.1029/2010GL044597.

Melzner, F., Gutowska, M. A., Langenbuch, M., Dupont, S., Lucassen, M., Thorndyke, M. C., et al. (2009). Physiological basis for high CO_2_ tolerance in marine ectothermic animals: pre-adaptation through lifestyle and ontogeny? Biogeosciences 6, 2313–2331. doi:https://doi.org/10.5194/bg-6-2313-2009.

Miller, K. J., Rowden, A. A., Williams, A., and Häussermann, V. (2011). Out of Their Depth? Isolated Deep Populations of the Cosmopolitan Coral *Desmophyllum dianthus* May Be Highly Vulnerable to Environmental Change. PLOS ONE 6, e19004. doi:10.1371/journal.pone.0019004.

Moheimani, N. R., Webb, J. P., and Borowitzka, M. A. (2012). Bioremediation and other potential applications of coccolithophorid algae: A review. Algal Res. 1, 120–133. doi:10.1016/j.algal.2012.06.002.

Moore, J. M., Carvajal, J. I., Rouse, G. W., and Wilson, N. G. (2018). The Antarctic Circumpolar Current isolates and connects: Structured circumpolarity in the sea star *Glabraster antarctica*. Ecol. Evol. 8, 10621–10633. doi:10.1002/ece3.4551.

Moore, K. M., Alderslade, P., and Miller, K. J. (2017). A taxonomic revision of *Anthothela* (Octocorallia: Scleraxonia: Anthothelidae) and related genera, with the addition of new taxa, using morphological and molecular data. Zootaxa 4304, 1–212. doi:10.11646/zootaxa.4304.1.1.

Moy, A. D., Howard, W. R., Bray, S. G., and Trull, T. W. (2009). Reduced calcification in modern Southern Ocean planktonic foraminifera. Nat. Geosci. 2, 276–280. doi:10.1038/ngeo460.

Müller, M. N., Trull, T. W., and Hallegraeff, G. M. (2017). Independence of nutrient limitation and carbon dioxide impacts on the Southern Ocean coccolithophore *Emiliania huxleyi*. ISME J. 11, 1777–1787. doi:10.1038/ismej.2017.53.

Müller, M., Trull, T., and Hallegraeff, G. (2015). Differing responses of three Southern Ocean *Emiliania huxleyi* ecotypes to changing seawater carbonate chemistry. Mar. Ecol. Prog. Ser. 531, 81–90. doi:10.3354/meps11309.

Nash, M. C., and Adey, W. (2017). Multiple phases of mg-calcite in crustose coralline algae suggest caution for temperature proxy and ocean acidification assessment: lessons from the ultrastructure and biomineralization in *Phymatolithon* (Rhodophyta, Corallinales). J. Phycol. 53, 970–984. doi:10.1111/jpy.12559.

Negrete-García, G., Lovenduski, N. S., Hauri, C., Krumhardt, K. M., and Lauvset, S. K. (2019). Sudden emergence of a shallow aragonite saturation horizon in the Southern Ocean. Nat. Clim. Change 9, 313–317. doi:10.1038/s41558-019-0418-8.

Nissen, C., Vogt, M., Münnich, M., Gruber, N., and Haumann, F. A. (2018). Factors controlling coccolithophore biogeography in the Southern Ocean. Biogeosciences 15, 6997–7024. doi:https://doi.org/10.5194/bg-15-6997-2018.

Orr, J. C., Fabry, V. J., Aumont, O., Bopp, L., Doney, S. C., Feely, R. a, et al. (2005). Anthropogenic ocean acidification over the twenty-first century and its impact on calcifying organisms. Nature 437, 681–686. doi:10.1038/nature04095.

Peck, L. S., Clark, M. S., Power, D., Reis, J., Batista, F. M., and Harper, E. M. (2015). Acidification effects on biofouling communities: winners and losers. Glob. Change Biol. 21, 1907–1913. doi:10.1111/gcb.12841.

Pierrat, B., Saucède, T., Laffont, R., Ridder, C. D., Festeau, A., and David, B. (2012). Large-scale distribution analysis of Antarctic echinoids using ecological niche modelling. Mar. Ecol. Prog. Ser. 463, 215–230. doi:10.3354/meps09842.

Post, A. L., O’Brien, P. E., Beaman, R. J., Riddle, M. J., and Santis, L. D. (2010). Physical controls on deep water coral communities on the George V Land slope, East Antarctica. Antarct. Sci. 22, 371–378. doi:10.1017/S0954102010000180.

Prather, C. M., Pelini, S. L., Laws, A., Rivest, E., Woltz, M., Bloch, C. P., et al. (2013). Invertebrates, ecosystem services and climate change. Biol. Rev. 88, 327–348. doi:10.1111/brv.12002.

Quartino, M. L., Deregibus, D., Campana, G. L., Latorre, G. E. J., and Momo, F. R. (2013). Evidence of Macroalgal Colonization on Newly Ice-Free Areas following Glacial Retreat in Potter Cove (South Shetland Islands), Antarctica. PLOS ONE 8, e58223. doi:10.1371/journal.pone.0058223.

Riebesell, U., Fabry, V., Hansson, L., and Gattuso, J. (2010). Guide to best practices for ocean acidification research and data reporting. Luxembourg: Publications Office of the European Union. Available at: https://op.europa.eu/en/publication-detail/-/publication/51887496-10b6-4b42-9876-1c2a1346fac5 [Accessed July 15, 2020].

Ries, J. B. (2011). A physicochemical framework for interpreting the biological calcification response to CO_2_-induced ocean acidification. Geochim. Cosmochim. Acta 75, 4053–4064. doi:10.1016/j.gca.2011.04.025.

Ries, J. B. (2012). A sea butterfly flaps its wings. Nat. Geosci. 5, 845–846. doi:10.1038/ngeo1655.

Ries, J. B., Cohen, A. L., and McCorkle, D. C. (2009). Marine calcifiers exhibit mixed responses to CO_2_-induced ocean acidification. Geology 37, 1131–1134. doi:10.1130/G30210A.1.

Rucker, J. B., and Carver, R. E. (1969). A Survey of the Carbonate Mineralogy of Cheilostome Bryozoa. J. Paleontol. 43, 791–799.

Schiebel, R. (2002). Planktic foraminiferal sedimentation and the marine calcite budget. Glob. Biogeochem. Cycles 16, 1–21. doi:10.1029/2001GB001459.

Schoenrock, K. M., Schram, J. B., Amsler, C. D., McClintock, J. B., Angus, R. A., and Vohra, Y. K. (2016). Climate change confers a potential advantage to fleshy Antarctic crustose macroalgae over calcified species. J. Exp. Mar. Biol. Ecol. 474, 58–66. doi:10.1016/j.jembe.2015.09.009.

Schram, J. B., Schoenrock, K. M., McClintock, J. B., Amsler, C. D., and Angus, R. A. (2016). Testing Antarctic resilience: the effects of elevated seawater temperature and decreased pH on two gastropod species. ICES J. Mar. Sci. 73, 739–752. doi:10.1093/icesjms/fsv233.

Seibel, B. A., and Dierssen, H. M. (2003). Cascading Trophic Impacts of Reduced Biomass in the Ross Sea, Antarctica: Just the Tip of the Iceberg? Biol. Bull. 205, 93–97. doi:10.2307/1543229.

Sewell, M. A., and Hofmann, G. E. (2011). Antarctic echinoids and climate change: a major impact on the brooding forms. Glob. Change Biol. 17, 734–744. doi:10.1111/j.1365-2486.2010.02288.x.

Sewell, M. A., Millar, R. B., Yu, P. C., Kapsenberg, L., and Hofmann, G. E. (2014). Ocean Acidification and Fertilization in the Antarctic Sea Urchin *Sterechinus neumayeri*: the Importance of Polyspermy. Environ. Sci. Technol. 48, 713–722. doi:10.1021/es402815s.

Smith, A. M., Berman, J., Key, M. M., and Winter, D. J. (2013a). Not all sponges will thrive in a high-CO_2_ ocean: Review of the mineralogy of calcifying sponges. Palaeogeogr. Palaeoclimatol. Palaeoecol. 392, 463–472. doi:10.1016/j.palaeo.2013.10.004.

Smith, A. M., Clark, D. E., Lamare, M. D., Winter, D. J., and Byrne, M. (2016). Risk and resilience: variations in magnesium in echinoid skeletal calcite. Mar. Ecol. Prog. Ser. 561, 1–16. doi:10.3354/meps11908.

Smith, A. M., Riedi, M. A., and Winter, D. J. (2013b). Temperate reefs in a changing ocean: Skeletal carbonate mineralogy of serpulids. Mar. Biol. 160, 2281–2294. doi:10.1007/s00227-013-2210-z.

Soest, R. W. M. V., Boury-Esnault, N., Vacelet, J., Dohrmann, M., Erpenbeck, D., Voogd, N. J. D., et al. (2012). Global Diversity of Sponges (Porifera). PLOS ONE 7, e35105. doi:10.1371/journal.pone.0035105.

Stark, J. S. (2008). Patterns of higher taxon colonisation and development in sessile marine benthic assemblages at Casey Station, Antarctica, and their use in environmental monitoring. Mar. Ecol. Prog. Ser. 365, 77–89. doi:10.3354/meps07559.

Stark, J. S., Roden, N. P., Johnstone, G. J., Milnes, M., Black, J. G., Whiteside, S., et al. (2018). Carbonate chemistry of an in-situ free-ocean CO_2_ enrichment experiment (antFOCE) in comparison to short term variation in Antarctic coastal waters. Sci. Rep. 8, 1–16. doi:10.1038/s41598-018-21029-1.

Steffel, B. V., Smith, K. E., Dickinson, G. H., Flannery, J. A., Baran, K. A., Rosen, M. N., et al. (2019). Characterization of the exoskeleton of the Antarctic king crab *Paralomis birsteini*. Invertebr. Biol. 138, e12246. doi:10.1111/ivb.12246.

Swezey, D. S., Bean, J. R., Ninokawa, A. T., Hill, T. M., Gaylord, B., and Sanford, E. (2017). Interactive effects of temperature, food and skeletal mineralogy mediate biological responses to ocean acidification in a widely distributed bryozoan. Proc. R. Soc. B Biol. Sci. 284, 20162349. doi:10.1098/rspb.2016.2349.

Tanur, A. E., Gunari, N., Sullan, R. M. A., Kavanagh, C. J., and Walker, G. C. (2010). Insights into the composition, morphology, and formation of the calcareous shell of the serpulid *Hydroides dianthus*. J. Struct. Biol. 169, 145–160. doi:10.1016/j.jsb.2009.09.008.

Taylor, M. L., and Rogers, A. D. (2015). Evolutionary dynamics of a common sub-Antarctic octocoral family. Mol. Phylogenet. Evol. 84, 185–204. doi:10.1016/j.ympev.2014.11.008.

Taylor, P. D., James, N. P., Bone, Y., Kuklinski, P., and Kyser, T. K. (2009). Evolving mineralogy of cheilostome bryozoans. Palaios 24, 440–452. doi:10.2110/palo.2008.p08.

Telesca, L., Peck, L. S., Sanders, T., Thyrring, J., Sejr, M. K., and Harper, E. M. (2019). Biomineralization plasticity and environmental heterogeneity predict geographical resilience patterns of foundation species to future change. Glob. Change Biol. 25, 4179–4193. doi:10.1111/gcb.14758.

Thatje, S. (2012). Effects of capability for dispersal on the evolution of diversity in Antarctic benthos. Integr. Comp. Biol. 52, 470–482. doi:10.1093/icb/ics105.

Thresher, R. E., Tilbrook, B., Fallon, S., Wilson, N. C., and Adkins, J. (2011). Effects of chronic low carbonate saturation levels on the distribution, growth and skeletal chemistry of deep-sea corals and other seamount megabenthos. Mar. Ecol. Prog. Ser. 442, 87–99. doi:10.3354/meps09400.

Thresher, R. E., Wilson, N. C., MacRae, C. M., and Neil, H. (2010). Temperature effects on the calcite skeletal composition of deep-water gorgonians (Isididae). Geochim. Cosmochim. Acta 74, 4655–4670. doi:10.1016/j.gca.2010.05.024.

Tittensor, D. P., Baco, A. R., Hall-Spencer, J. M., Orr, J. C., and Rogers, A. D. (2010). Seamounts as refugia from ocean acidification for cold-water stony corals. Mar. Ecol. 31, 212–225. doi:10.1111/j.1439-0485.2010.00393.x.

Turley, C. M., Roberts, J. M., and Guinotte, J. M. (2007). Corals in deep-water: will the unseen hand of ocean acidification destroy cold-water ecosystems? Coral Reefs 26, 445–448. doi:10.1007/s00338-007-0247-5.

Vaughan, D., and Marshall, G. (2003). Recent Rapid Regional Climate Warming on the Antarctic Peninsula. Clim. Change, 243–274.

Waldbusser, G. G., Hales, B., Langdon, C. J., Haley, B. A., Schrader, P., Brunner, E. L., et al. (2015). Saturation-state sensitivity of marine bivalve larvae to ocean acidification. Nat. Clim. Change 5, 273–280. doi:10.1038/nclimate2479.

Waller, C. L., Overall, A., Fitzcharles, E. M., and Griffiths, H. (2017). First report of *Laternula elliptica* in the Antarctic intertidal zone. Polar Biol. 40, 227–230. doi:10.1007/s00300-016-1941-y.

Watling, L., France, S. C., Pante, E., and Simpson, A. (2011). “Chapter Two - Biology of Deep-Water Octocorals,” in Advances in Marine Biology Advances in Marine Biology., ed. M. Lesser (Academic Press), 41–122. doi:10.1016/B978-0-12-385529-9.00002-0.

Watson, S.-A., Peck, L. S., Tyler, P. A., Southgate, P. C., Tan, K. S., Day, R. W., et al. (2012). Marine invertebrate skeleton size varies with latitude, temperature and carbonate saturation: implications for global change and ocean acidification. Glob. Change Biol. 18, 3026–3038. doi:10.1111/j.1365-2486.2012.02755.x.

Weiner, S., and Dove, P. M. (2003). An overview of biomineralization processes and the problem of the vital effect. Rev. Mineral. Geochem. 54, 1–29. doi:10.2113/0540001.

Weyl, P. K. (1959). The change in solubility of calcium carbonate with temperature and carbon dioxide content. Geochim. Cosmochim. Acta 17, 214–225. doi:10.1016/0016-7037(59)90096-1.

Wiencke, C., Amsler, C. D., and Clayton, M. N. (2014). Macroalgae. Biogeogr. Atlas South. Ocean. Available at: http://data.biodiversity.aq/ [Accessed February 8, 2020].

Wright, J. T., and Gribben, P. E. (2017). Disturbance-mediated facilitation by an intertidal ecosystem engineer. Ecology 98, 2425–2436. doi:10.1002/ecy.1932.

Yu, P. C., Sewell, M. A., Matson, P. G., Rivest, E. B., Kapsenberg, L., and Hofmann, G. E. (2013). Growth Attenuation with Developmental Schedule Progression in Embryos and Early Larvae of *Sterechinus neumayeri* Raised under Elevated CO_2_. PLOS ONE 8, e52448. doi:10.1371/journal.pone.0052448.

Zeebe, R. E. (2012). History of Seawater Carbonate Chemistry, Atmospheric CO_2_, and Ocean Acidification. Annu. Rev. Earth Planet. Sci. 40, 141–165. doi:10.1146/annurev-earth-042711-105521.

