## Supplemental files for "Predicting potential impacts of ocean acidification on marine calcifiers from the Southern Ocean"

### *Supplementary Material*

#### Supplementary Figures and Tables

**Figure S1.** Forest plots of all studies on the effect of altered carbonate chemistry on marine calcifiers south of 60ºS included in the meta-analysis. The data is separated by mineralogical composition and information is provided on the study paper, experiment (where applicable) and species investigated. For each study, the response ratio, variance (v) and 95% confidence interval is shown. At the end of each mineralogical composition bracket, summary statistics from weighted, random effects are provided including the Q statistic, degrees of freedom, p-value, $I^2$, mean response ratio and 95% confidence internal.

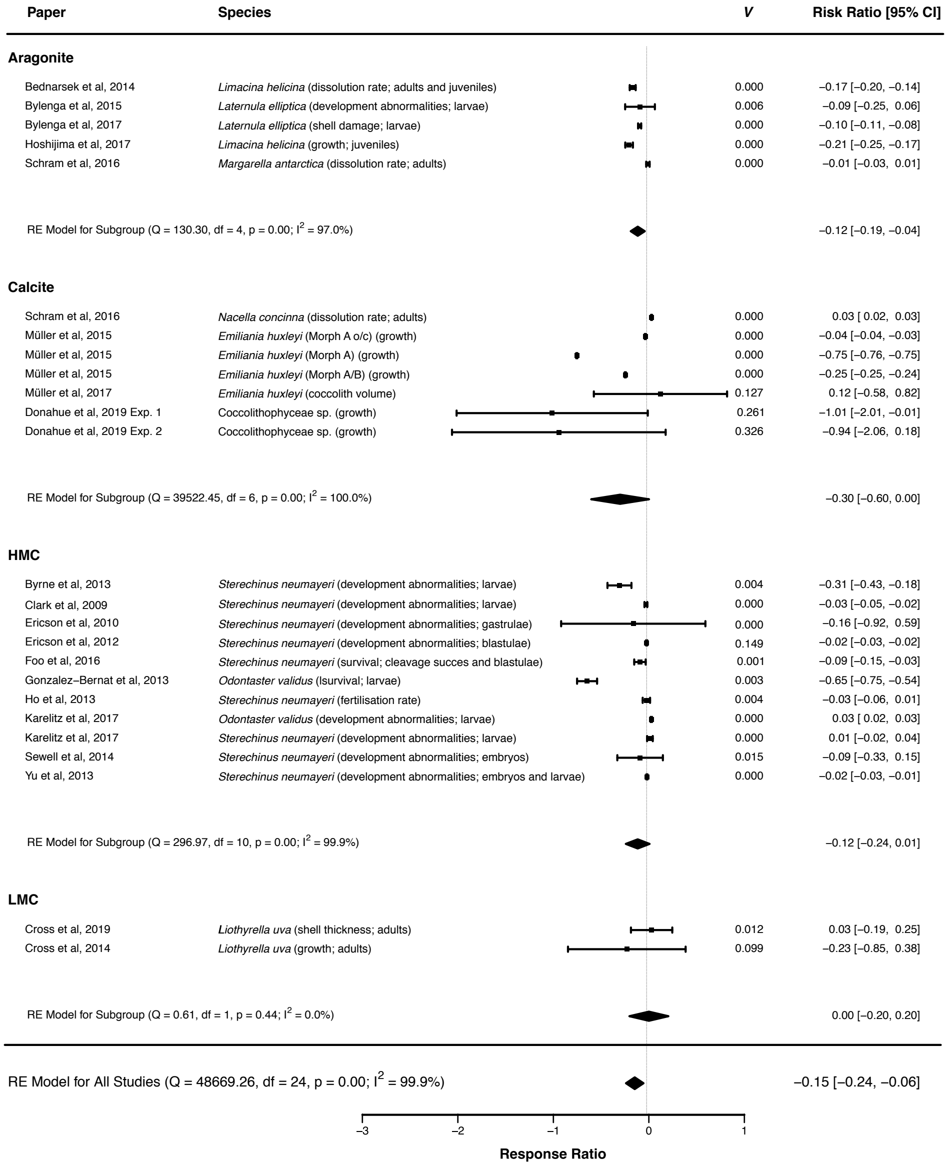

**Table S1.** Mean percentages of $MgCO_3$ in calcite by weight in skeletons of Antarctic echinoderms using data from the literature (echinoids (test only), holothuroids (calcareous ring), ophiuroids (arm and disk), and asteroids (arm or intact)). LMC: low-Mg calcite (0–4 wt% $MgCO_3$); IMC: intermediate Mg calcite (4–8 wt% $MgCO_3$); HMC: high Mg calcite (> 8 wt% $MgCO_3$).

| Species | Collection location | Latitude (S) | Longitude (W) | Depth (m) | Mean wt% $MgCO_3$ in calcite | Standard error | n | Category | Reference |
|---|---|---|---|---|---|---|---|---|---|
| *Acodontaster conspicuus* (Koehler, 1920) | Lemaire Channel | 65º04.66' | 63º58.21' | 5–40 | 9.39 | | 1 | HMC | McClintock et al. 2011 |
| *Acodontaster hodgsoni* (Bell, 1908) | Dallmann Bay | 64º09.45' | 62º44.73' | 150–170 | 9.85 | 0.02 | 3 | HMC | McClintock et al. 2011 |
| *Amphipneustes similis* Mortensen, 1936 | Hugo Island | 64º45.53' | 64º28.26' | 670–700 | 7.51 | 0.74 | 4 | HMC | McClintock et al. 2011 |
| *Bathybiaster loripes* Sladen, 1889 | Western Antarctic Peninsula | - | - | 341–764 | 6.09 | 0.004 | 16 | IMC | Duquette et al. 2018 |
| *Ctenocidaris perrieri* Koehler, 1912 | Hugo Island | 64º45.53' | 64º28.26' | 670–700 | 7.61 | 0.24 | 3 | HMC | McClintock et al. 2011 |
| *Diplasterias brandti* (Bell, 1881) | Arthur Harbor | 64º46.47' | 64º03.29' | 5–40 | 9.52 | 0.15 | 3 | HMC | McClintock et al. 2011 |
| *Diplopteraster verrucosus* (Sladen, 1882) | Banana Trench | 66º17.63' | 66º36.18' | 850–950 | 8.12 | 0.01 | 2 | HMC | McClintock et al. 2011 |
| *Glabraster antarctica* (E. A. Smith, 1876) | Dallmann Bay | 64º09.45' | 62º44.73' | 150–170 | 10.2, | 0.07 | 3 | HMC, | McClintock et al. 2011 |
| | Western Antarctic Peninsula | - | - | 341–764 | 7.84 | 0.006 | 10 | IMC | Duquette et al. 2018 |
| *Granaster nutrix* (Studer, 1885) | Dallmann Bay | 64º09.45' | 62º44.73' | 150–170 | 8.29 | 0.38 | 5 | HMC | McClintock et al. 2011 |
| *Kampylaster incurvatus* Koehler, 1920 | Elephant Island | 61º12.81' | 56º01.11' | 160–170 | 9.28 | 0.63 | 4 | HMC | McClintock et al. 2011 |
| *Labidiaster annulatus* Sladen, 1889 | Low Island | 63º31.84' | 62º45.07' | 140–215 | 9.79 | 0.04 | 9 | HMC | McClintock et al. 2011 |
| *Macroptychaster accrescens* (Koehler, 1920) | Hugo Island | 64º45.53' | 64º28.26' | 670–700 | 9.83 | 0.16 | 3 | HMC, | McClintock et al. 2011 |
| | Western Antarctic Peninsula | - | - | 341–764 | 6.09 | 0.010 | 2 | IMC | Duquette et al. 2018 |

| | | | | | | | | | |
|---|---|---|---|---|---|---|---|---|---|
| *Molpadia musculus* Risso, 1826 | Arthur Harbor | 64º46.47' | 66º03.29' | 5–40 | 8.26 | 0.17 | 3 | HMC | McClintock et al. 2011 |
| *Neosmilaster georgianus* (Studer, 1885) | SE Bonaparte Pt. | 64º46.47' | 64º02.53' | 5–40 | 9.37 | 0.05 | 3 | HMC | McClintock et al. 2011 |
| *Odontaster meridionalis* (E. A. Smith, 1876) | Dallmann Bay | 64º09.45' | 62º44.73' | 150–170 | 9.5 | 0.04 | 3 | HMC | McClintock et al. 2011 |
| *Odontaster penicillatus* (Philippi, 1870) | Dallmann Bay | 64º09.45' | 62º44.73' | 150–170 | 9.91 | | 1 | HMC | McClintock et al. 2011 |
| *Ophiacantha antarctica* Koehler, 1900 | Western Antarctic Peninsula | - | - | 341–764 | 8.48 | 0.004 | 57 | HMC | Duquette et al. 2018 |
| *Ophiocten megaloplax* Koehler, 1900 | Western Antarctic Peninsula | - | - | 341–764 | 3.81 | 0.005 | 4 | LMC | Duquette et al. 2018 |
| *Ophiolimna antarctica* (Lyman, 1879) | Western Antarctic Peninsula | - | - | 341–764 | 7.26 | 0.005 | 35 | IMC | Duquette et al. 2018 |
| *Ophionotus victoriae* Bell, 1902 | Arthur Harbor | 64º46.47' | 66º03.29' | 5–40 | 9.2 | 0.1 | 3 | HMC | McClintock et al. 2011 |
| | Western Antarctic Peninsula | - | - | 341–764 | 7.33 | 0.007 | 10 | IMC | Duquette et al. 2018 |
| *Ophioperla koehleri* (Bell, 1908) | Western Antarctic Peninsula | - | - | 341–764 | 6.6 | 0.005 | 20 | IMC | Duquette et al. 2018 |
| *Ophiosparte gigas* Koehler, 1922 | Arthur Harbor | 64º46.47' | 66º03.29' | 5–40 | 9.13 | | 1 | HMC | McClintock et al. 2011 |
| *Ophiura (Ophiuroglypha) carinifera* (Koehler, 1901) | Western Antarctic Peninsula | - | - | 341–764 | 7.33 | 0.006 | 17 | IMC | Duquette et al. 2018 |
| *Paralophaster godfroyi* (Koehler, 1912) | Stepping Stones Is. | 64º47.18' | 63º59.85' | 5–40 | 9.59 | | 1 | HMC | McClintock et al. 2011 |
| *Perknaster aurorae* (Koehler, 1920) | Low Island | 63º31.84' | 62º45.07' | 140–215 | 8.86 | 0.55 | 3 | HMC | McClintock et al. 2011 |
| *Perknaster densus* Sladen, 1889 | Lemaire Channel | 65º04.66' | 63º58.21' | 5–40 | 7.47 | | 1 | HMC | McClintock et al. 2011 |
| *Perknaster fuscus* Sladen, 1889 | Dallmann Bay | 64º09.45' | 62º44.73' | 150–170 | 9.92 | | 1 | HMC | McClintock et al. 2011 |
| *Pseudostichopus spiculiferus* (O'Loughlin, 2002) | Banana Trench | 66º17.63' | 66º36.18' | 850–950 | 7.49 | | 1 | HMC | McClintock et al. 2011 |

| | | | | | | | | | |
|---|---|---|---|---|---|---|---|---|---|
| *Sterechinus neumayeri* (Meissner, 1900) | Hugo Island | 64º45.53' | 65º28.26' | 670–700 | 6.04 | 0.1 | 6 | HMC | McClintock et al. 2011 |
| | Norsel Point | 64º45.62' | 64º05.90' | 5–40 | | | | | |
| | Lemaire Channel | 65º04.66' | 63º58.21' | 5–40 | | | | | |

**Table S2.** Mean percentages of $MgCO_3$ in calcite by weight in skeletons of Antarctic bryozoans using data from the literature. A = Antarctica; BB = Brown Bay; BBy = Borge Bay; CB = central basin; IEI = inner Ezcurra Inlet; KG= King George Island; OB = O'Brien Bay; OEI = outer Ezcurra Inlet; OO= Off Oates Land; MM = McMurdo Bay; PA = Palmer Archipelago; RB = Ryder Bay; RS = Ross Sea; SA = Scotia Arc; SB = Shannon Bay; SS= South Shetland Islands; Terre Adélie = TA WQ = Winter Quarters Bay. LMC = low-Mg calcite (0–4 wt% $MgCO_3$); IMC = intermediate Mg calcite (4–8 wt% $MgCO_3$); HMC = high Mg calcite (> 8 wt% $MgCO_3$).

| **Species** | **Collection location** | **Latitude (S)** | **Longitude (W)** | **Depth (m)** | **Mean wt% $MgCO_3$ in calcite** | **Standard deviation** | **n** | **Category** | **Reference** |
|---|---|---|---|---|---|---|---|---|---|
| *Aimulosia australis* Jullien, 1888 | OEI | 62°09′ | 52°31′ | 6–15 | 3.91 | 1.98 | 4 | LMC | Krzeminska et al. 2016 |
| *Amastigia gaussi* (Kluge 1914) | CB | 62°09′ | 52°27′ | 115 | 2.37 | 0.00 | 2 | LMC | Krzeminska et al. 2016 |
| *Amphiblestrum inermis* (Kluge 1914) | OEI | 62°09′ | 52°31′ | 6–15 | 5.59 | 0.52 | 7 | IMC | Krzeminska et al. 2016 |
| *Antarcticaetos bubeccata* (Rogick 1955) | RS | 76º | - | - | 3.9 | - | 1 | LMC | Taylor et al. 2009 |
| | OEI | 62°09′ | 52°31′ | 110–117 | 5.82 | 1.67 | 3 | IMC | Krzeminska et al. 2016 |
| | CB | 62°09′ | 52°27′ | 60–251 | 3.60 | 0.46 | 5 | LMC | Krzeminska et al. 2016 |
| *Antarctothoa antarctica* Moyano & Gordon, 1980 | RB | 68º | 68º | 8–9 | 0.6 | 0.531 | 159 | LMC | Loxton et al. 2014 |
| | IEI | 62°10′ | 58°35′ | 85 | 0.67 | - | 1 | LMC | Krzeminska et al. 2016 |
| *Antarctothoa bougainvillei* (d'Orbigny 1842) | OEI | 62°09′ | 52°31′ | 6–15 | 1.27 | 0.62 | 9 | LMC | Krzeminska et al. 2016 |
| | CB | 62°09′ | 52°27′ | 70 | 1.9 | - | 1 | LMC | |
| *Arachnopusia columnaris* Hayward & Thorpe, 1988 | PA | 64º | - | - | 3.9 | - | 1 | LMC | Taylor et al. 2009 |
| | OEI | 62°09′ | 52°31′ | 6–15 | 5.9 | 0.25 | 6 | IMC | Krzeminska et al. 2016 |
| | OEI | 62°09′ | 52°31′ | 105 | 5.35 | - | 1 | IMC | Krzeminska et al. 2016 |
| | CB | 62°09′ | 52°27′ | 70 | 4.67 | - | 1 | IMC | Krzeminska et al. 2016 |

| | | | | | | | | | |
|---|---|---|---|---|---|---|---|---|---|
| *Arachnopusia decipiens* Hayward & Thorpe, 1988 | OB | 66°17′ | - | 6–22 | 6.30 | 0.74 | 53 | IMC | Figuerola et al. 2019 |
| *Astochoporella cassidula* Hayward & Thorpe, 1988 | RS | 76º | - | - | 4.84 | - | 1 | IMC | Taylor et al. 2009 |
| *Austroflustra vulgaris* (Kluge 1914) | A | - | - | - | 5 | - | 1 | IMC | Borisenko & Gontar 1991 |
| | CB | 62°09′ | 52°27′ | 60 | 4.42 | 0.73 | 2 | IMC | Krzeminska et al. 2016 |
| *Beania erecta* Waters, 1904 | OEI | 62°09′ | 52°31′ | 70 | 7.17 | 0.86 | 2 | IMC | Krzeminska et al. 2016 |
| | OEI | 62°09′ | 52°31′ | 6–10 | 5.75 | 0.18 | 2 | IMC | Krzeminska et al. 2016 |
| | CB | 62°09′ | 52°27′ | 60 | 4.67 | - | 1 | IMC | Krzeminska et al. 2016 |
| | SB | 66°16' | - | 6–22 | 7.80 | 1.20 | 28 | IMC | Figuerola et al. 2019 |
| | BB | 66°16' | - | 6–19 | 7.51 | 1.37 | 28 | IMC | Figuerola et al. 2019 |
| | OB | 66°17′ | - | 8–22 | 8.58 | 1.68 | 47 | IMC | Figuerola et al. 2019 |
| *Beania livingstonei* Hastings, 1943 | A | - | - | - | 4.00 | - | 1 | IMC | Borisenko & Gontar 1991 |
| *Bostrychopora dentata* (Waters, 1904) | RS | 75º | - | - | 4.11 | - | 1 | IMC | Taylor et al. 2009 |
| *Brettiopsis triplex* (Hastings 1943) | CB | 62°09′ | 52°27′ | 104 | 6.13 | - | 1 | IMC | Krzeminska et al. 2016 |
| *Caberea darwinii* Busk, 1884 | CB | 62°09′ | 52°27′ | 70 | 4.16 | - | 1 | IMC | Krzeminska et al. 2016 |
| | OEI | 62°09′ | 52°31′ | 109–130 | 5.36 | 0.32 | 4 | IMC | Krzeminska et al. 2016 |
| | IEI | 62°10′ | 58°35′ | 113–134 | 3.90 | 0.79 | 5 | LMC | Krzeminska et al. 2016 |
| *Camptoplites angustus* (Kluge, 1914) | A | - | - | - | 4.00 | - | 1 | IMC | Borisenko & Gontar 1991 |
| *Camptoplites bicornis* (Busk 1884) | CB | 62°09′ | 52°27′ | 251 | 3.53 | - | 1 | LMC | Krzeminska et al. 2016 |
| *Camptoplites* cf. *angustus* (Kluge 1914) | IEI | 62°10′ | 58°35′ | 119 | 3.09 | - | 1 | LMC | Krzeminska et al. 2016 |

| | | | | | | | | | |
|---|---|---|---|---|---|---|---|---|---|
| *Camptoplites latus* (Kluge 1914) | CB | 62°09′ | 52°27′ | 251 | 2.44 | - | 1 | LMC | Krzeminska et al. 2016 |
| *Camptoplites retiformis* (Kluge 1914) | CB | 62°09′ | 52°27′ | 296 | 1.94 | 1.97 | 1 | LMC | Krzeminska et al. 2016 |
| | CB | 62°09′ | 52°27′ | 70 | 5.35 | 0.00 | 2 | IMC | Krzeminska et al. 2016 |
| *Camptoplites tricornis* (Waters 1904) | IEI | 62°10′ | 58°35′ | 85 | 3.13 | 0.00 | 2 | LMC | Krzeminska et al. 2016 |
| *Carbasea curva* (Kluge 1914) | A | - | - | - | 5 | - | 1 | IMC | Borisenko & Gontar 1991 |
| | CB | 62°09′ | 52°27′ | 115 | 7.26 | 0.86 | 2 | IMC | Krzeminska et al. 2016 |
| *Carbasea ovoidea* Busk, 1852 | CB | 62°09′ | 52°27′ | 60 | 3.64 | - | 1 | LMC | Krzeminska et al. 2016 |
| | CB | 62°09′ | 52°27′ | 232 | 3.99 | 1.04 | 3 | LMC | Krzeminska et al. 2016 |
| *Cellaria aurorae* Livingstone, 1928 | CB | 62°09′ | 52°27′ | 245 | 2.37 | - | 1 | LMC | Krzeminska et al. 2016 |
| *Cellaria diversa* Livingstone, 1928 | IEI | 62°10′ | 58°35′ | 72 | 2.37 | - | 1 | LMC | Krzeminska et al. 2016 |
| | OEI | 62°09′ | 52°31′ | 107–123 | 2.84 | 0.50 | 3 | LMC | Krzeminska et al. 2016 |
| | CB | 62°09′ | 52°27′ | 60 | 1.27 | 0.29 | 3 | | Krzeminska et al. 2016 |
| | CB | 62°09′ | 52°27′ | 115–205 | 2.26 | 0.60 | 3 | LMC | Krzeminska et al. 2016 |
| *Cellarinella latilaminata* Moyano, 1974 | IEI | 62°10′ | 58°35′ | 113 | 3.64 | - | 1 | LMC | Krzeminska et al. 2016 |
| *Cellarinella margueritae* Rogick, 1956 | TA | 66º | | 180–346 | 4.6 | - | 4 | LMC | Loxton et al. 2013 |
| | CB | 62°09′ | 52°27′ | 70 | 3.14 | 1.08 | 2 | LMC | Krzeminska et al. 2016 |
| *Cellarinella nodulata* Waters, 1904 | CB | 62°09′ | 52°27′ | 270–290 | 3.20 | 0.38 | 4 | LMC | Krzeminska et al. 2016 |
| *Cellarinella nutti* Rogick, 1956 | CB | 62°09′ | 52°27′ | 115 | 4.05 | 1.60 | 3 | IMC | Krzeminska et al. 2016 |
| *Cellarinella rogickae* Moyano, 1965 | CB | 62°09′ | 52°27′ | 115 | 5.87 | - | 1 | IMC | Krzeminska et al. 2016 |

| | | | | | | | | |
|---|---|---|---|---|---|---|---|---|
| *Cellarinella terminata* Hayward & Winston, 1994 | CB | 62°09′ | 52°27′ | 104 | 3.65 | - | 1 | LMC | Krzeminska et al. 2016 |
| *Cellarinella virgula* Hayward & Ryland, 1991 | IEI | 62°10′ | 58°35′ | 85 | 4.16 | - | 1 | IMC | Krzeminska et al. 2016 |
| *Cellarinelloides crassus* Moyano, 1970 | RS | 76º | - | - | 4.84 | - | 1 | IMC | Taylor et al. 2009 |
| | TA | 66°53′ | - | 347 | 5.19 | - | 3 | IMC | Loxton et al. 2013 |
| *Chaperiopsis signyensis* Hayward, 1993 | OEI | 62°09′ | 52°31′ | 10–15 | 4.76 | 0.30 | 3 | IMC | Krzeminska et al. 2016 |
| *Chondriovelum adeliense* (Livingstone 1928) | OO | 70º | - | - | 4.50 | - | 1 | IMC | Taylor et al. 2009 |
| | CB | 62°09′ | 52°27′ | 333 | 3.64 | - | 1 | LMC | Krzeminska et al. 2016 |
| *Diastopora solida* Waters, 1904 | A | - | - | - | 5.00 | - | 1 | IMC | Borisenko & Gontar 1991 |
| *Ellisina antarctica* Hastings, 1945 | OEI | 62°09′ | 52°31′ | 6–15 | 5.63 | 1.66 | 8 | IMC | Krzeminska et al. 2016 |
| | BB | 66°16' | - | 6–19 | 6.23 | 0.49 | 13 | IMC | Figuerola et al. 2019 |
| | OB | 66°17′ | - | 6–22 | 5.80 | 0.65 | 13 | ICM | Figuerola et al. 2019 |
| *Eminooecia carsonae* (Rogick, 1957) | WQ | 77º | - | - | 4.84 | - | 1 | IMC | Taylor et al. 2009 |
| *Escharoides tridens* (Calvet 1909) | OEI | 62°09′ | 52°31′ | 6–15 | 2.94 | 0.65 | 6 | LMC | Krzeminska et al. 2016 |
| *Exochella elegans* Hayward, 1991 | SC | 60º | - | - | 6.13 | - | 1 | IMC | Taylor et al. 2009 |
| *Fasciculipora ramosa* d'Orbigny, 1842 | A | - | - | - | 5.0 | - | 1 | IMC | Borisenko & Gontar 1991 |
| | TA | 66º | - | 185–598 | 3.9 | 053 | 32 | LMC | Figuerola et al. 2015 |
| *Favosthimosia milleporoides* (Calvet 1909) | OEI | 62°09′ | 52°31′ | 110 | 3.65 | - | 1 | LMC | Krzeminska et al. 2016 |
| *Fenestrulina rugula* Hayward & Ryland, 1990 | RB | 68º | 68º | 8–9 | 4.92 | 0.77 | 190 | IMC | Loxton et al. 2014 |
| | OEI | 62°09′ | 52°31′ | 6 | 5.33 | 0.89 | 7 | IMC | Krzeminska et al. 2016 |

| | | | | | | | | | |
|---|---|---|---|---|---|---|---|---|---|
| *Filaguria spatulata* (Calvet 1909) | OEI | 62°09′ | 52°31′ | 6–15 | 5.34 | 0.63 | 4 | IMC | Krzeminska et al. 2016 |
| | OEI | 62°09′ | 52°31′ | 108 | 1.94 | - | 1 | LMC | Krzeminska et al. 2016 |
| *Himantozoum (Himantozoum) antarcticum* (Calvet 1909) | A | - | - | - | 4 | - | 1 | IMC | Borisenko & Gontar 1991 |
| | CB | 62°09′ | 52°27′ | 46 | 3.90 | 1.09 | 2 | LMC | Krzeminska et al. 2016 |
| *Hippadenella inerma* (Calvet, 1909) | RB | 68º | 68º | 8–9 | 5.02 | 0.68 | 145 | IMC | Loxton et al. 2014 |
| *Inversiula nutrix* Jullien, 1888 | KG | 60º | - | - | 0.9 | - | 1 | LMC | Taylor et al. 2009 |
| | SS | 60º | - | - | 3.1 | - | 1 | LMC | Taylor et al. 2009 |
| | RB | 68º | 68º | 8–9 | 2.64 | 0.26 | 90 | LMC | Loxton et al. 2014 |
| | OEI | 62°09′ | 52°31′ | 6 | 3.80 | 1.22 | 8 | LMC | Krzeminska et al. 2016 |
| | BB | 66°16' | - | 6–19 | 2.00 | 1.08 | 59 | LMC | Figuerola et al. 2019 |
| | OB | | - | 8–22 | 2.18 | 0.93 | 39 | LMC | Figuerola et al. 2019 |
| | SB | 66°16' | - | 6–22 | 1.74 | 0.72 | 27 | LMC | Figuerola et al. 2019 |
| *Isoschizoporella tricuspis* (Calvet 1909) | CB | 62°09′ | 52°27′ | 6–19 | 6.54 | 1.26 | 3 | IMC | Krzeminska et al. 2016 |
| *Isosecuriflustra angusta* (Kluge 1914) | TA | 66º | - | 187–262 | 4.83 | - | 3 | IMC | Loxton et al. 2013 |
| | OEI | 62°09′ | 52°31′ | 108 | 4.41 | | 1 | IMC | Krzeminska et al. 2016 |
| | IEI | 62°10′ | 58°35′ | 93–134 | 5.46 | 0.74 | 7 | IMC | Krzeminska et al. 2016 |
| *Isosecuriflustra tenuis* (Kluge 1914) | IEI | 62°10′ | 58°35′ | 73–120 | 5.70 | 1.03 | 7 | IMC | Krzeminska et al. 2016 |
| *Isosecuriflustra thysanica* (Moyano 1972) | OEI | 62°09′ | 52°31′ | 113 | 3.39 | - | 1 | LMC | Krzeminska et al. 2016 |

| | | | | | | | | | |
|---|---|---|---|---|---|---|---|---|---|
| *Klugeflustra antarctica* (Hastings 1943) | CB | 62°09′ | 52°27′ | 290 | 5.10 | 1.46 | 2 | IMC | Krzeminska et al. 2016 |
| *Klugeflustra drygalskii* (Kluge 1914) | IEI | 62°10′ | 58°35′ | 115 | 3.13 | - | 1 | LMC | Krzeminska et al. 2016 |
| *Klugeflustra vanhoeffeni* (Kluge 1914) | CB | 62°09′ | 52°27′ | 70 | 5.87 | 0.26 | 3 | IMC | Krzeminska et al. 2016 |
| *Kymella polaris* (Waters 1904) | RS | 76º | - | - | 4.50 | - | 1 | IMC | Taylor et al. 2009 |
| | CB | 62°09′ | 52°27′ | 70–115 | 5.30 | 0.75 | 3 | IMC | Krzeminska et al. 2016 |
| *Lacerna eatoni* (Busk 1876) | OEI | 62°09′ | 52°31′ | 6–15 | 5.46 | 0.84 | 7 | IMC | Krzeminska et al. 2016 |
| *Lacerna hosteensis* Jullien, 1888 | CB | 62°09′ | 52°27′ | 60 | 3.90 | - | 1 | LMC | Krzeminska et al. 2016 |
| *Lageneschara lyrulata* (Calvet, 1909) | MM | 78º | - | - | 3.99 | - | 1 | LMC | Taylor et al. 2009 |
| | BBy | 59º | - | - | 3.99 | 0.73 | 2 | | Taylor et al. 2009 |
| | TA | 66º | - | 185–598 | 4.0 | 0.39 | 32 | LMC | Figuerola et al. 2015 |
| *Larvapora mawsoni* (Livingstone, 1928) | PA | 65º | - | - | 4.48 | - | 1 | IMC | Taylor et al. 2009 |
| *Melicerita* cf. *flabellifera* Hayward & Winston, 1994 | IEI | 62°10′ | 58°35′ | 120 | 3.51 | 2.29 | 3 | LMC | Krzeminska et al. 2016 |
| | CB | 62°09′ | 52°27′ | 232 | 3.52 | 0.91 | 2 | LMC | Krzeminska et al. 2016 |
| *Melicerita obliqua* (Thornely, 1924) | A | - | - | - | 5.0 | - | 1 | IMC | Sandberg 1977, Borisenko & Gontar 1991 |
| | TA | 66º | - | 385–598 | 4.2 | 0.82 | 10 | IMC | Figuerola et al. 2015 |
| *Micropora notialis* Hayward & Ryland, 1993 | OEI | 62°09′ | 52°31′ | 6–15 | 6.14 | 1.10 | 8 | IMC | Krzeminska et al. 2016 |
| *Microporella stenoporta* Hayward & Taylor, 1984 | OEI | 62°09′ | 52°31′ | 15 | 5.22 | - | 1 | IMC | Krzeminska et al. 2016 |
| *Nematoflustra flagellata* (Waters 1904) | A | - | - | 5 | - | - | 1 | LMC | Borisenko & Gontar 1991 |

| | | | | | | | | | |
|---|---|---|---|---|---|---|---|---|---|
| | OEI | 62°09′ | 52°31′ | 60 | 1.70 | - | 1 | LMC | Krzeminska et al. 2016 |
| | OEI | 62°09′ | 52°31′ | 109–272 | 3.10 | 1.68 | 4 | LMC | Krzeminska et al. 2016 |
| *Notoplites drygalskii* (Kluge 1914) | CB | 62°09′ | 52°27′ | 251 | 2.54 | 0.12 | 2 | LMC | Krzeminska et al. 2016 |
| | IEI | 62°10′ | 58°35′ | 85–134 | 3.58 | 0.97 | 4 | LMC | Krzeminska et al. 2016 |
| *Notoplites tenuis* (Kluge 1914) | CB | 62°09′ | 52°27′ | 70 | 6.13 | - | 1 | IMC | Krzeminska et al. 2016 |
| | CB | 62°09′ | 52°27′ | 251 | 5.10 | - | 1 | IMC | Krzeminska et al. 2016 |
| *Orthoporidra compacta* (Waters 1904) | PA | 64º | - | - | 5.10 | - | 1 | IMC | Taylor et al. 2009 |
| | IEI | 62°10′ | 58°35′ | 114–115 | 7.00 | 1.96 | 2 | IMC | Krzeminska et al. 2016 |
| *Osthimosia clavata* Waters, 1904 | A | - | - | - | 7.00 | - | 1 | IMC | Borisenko & Gontar 1991 |
| *Osthimosia* cf. *malingae* Hayward, 1992 | OEI | 62°09′ | 52°31′ | 117–123 | 5.53 | 1.95 | 2 | IMC | Krzeminska et al. 2016 |
| *Osthimosia* cf. *curtioscula* Hayward, 1992 | CB | 62°09′ | 52°27′ | 60–70 | 4.76 | 0.40 | 4 | IMC | Krzeminska et al. 2016 |
| *Osthimosia mariae* Hayward, 1992 | IEI | 62°10′ | 58°35′ | 120 | 7.17 | - | 1 | IMC | Krzeminska et al. 2016 |
| *Osthimosia notialis* Hayward, 1992 | CB | 62°09′ | 52°27′ | 232–251 | 5.56 | 1.19 | 3 | IMC | Krzeminska et al. 2016 |
| *Paracellaria wandeli* (Calvet, 1909) | PA | 65º | - | - | 2.71 | - | 1 | LMC | Taylor et al. 2009 |
| *Pemmatoporella marginata* (Calvet, 1909) | RS | 76º | - | - | 5.27 | - | 1 | IMC | Taylor et al. 2009 |
| *Polirhabdotos inclusum* (Waters, 1904) | PA | 65º | - | - | 4.59 | 0.97 | 2 | IMC | Taylor et al. 2009 |
| *Reteporella frigida* (Waters 1904) | IEI | 62°10′ | 58°35′ | 115–122 | 6.83 | 2.47 | 4 | IMC | Krzeminska et al. 2016 |
| *Reteporella hippocrepis* (Waters 1904) | A | - | - | - | 7.00 | - | 1 | IMC | Borisenko & Gontar 1991 |
| *Rhynchozoon fistulosum* Hayward, 1993 | CB | 62°09′ | 52°27′ | 252 | 4.33 | 0.30 | 3 | IMC | Krzeminska et al. 2016 |

| | | | | | | | | | |
|---|---|---|---|---|---|---|---|---|---|
| *Smittina alticollarita* Rogick, 1956 | CB | 62°09′ | 52°27′ | 70 | 5.62 | - | 1 | IMC | Krzeminska et al. 2016 |
| *Smittina antarctica* (Waters 1904) | RS | 76º | - | - | 5.7 | - | 1 | - | Taylor et al. 2009 |
| | CB | 62°09′ | 52°27′ | 251 | 6.18 | 1.56 | 3 | IMC | Krzeminska et al. 2016 |
| *Smittina directa* (Waters 1904) | RS | 75º | - | - | 4.50 | - | 1 | IMC | Taylor et al. 2009 |
| *Smittina obicullata* Rogick, 1956 | CB | 62°09′ | 52°27′ | 251 | 5.10 | - | 1 | IMC | Krzeminska et al. 2016 |
| *Smittina pocilla* Hayward & Thorpe, 1990 | OEI | 62°09′ | 52°31′ | 6–15 | 6.28 | 1.82 | 3 | IMC | Krzeminska et al. 2016 |
| *Swanomia membranacea* (Thornely 1924) | IEI | 62°10′ | 58°35′ | 113 | 5.06 | 2.25 | 2 | IMC | Taylor et al. 2009 |
| | OEI | 62°09′ | 52°31′ | 122 | 4.67 | - | 1 | IMC | Krzeminska et al. 2016 |
| *Systenopora contracta* Waters, 1904 | RS | 75º | - | - | 3.02 | 1.16 | 2 | LMC | Taylor et al. 2009 |
| | TA | 66º | | 443–660 | 3.8 | 0.42 | 30 | LMC | Figuerola et al. 2015 |
| *Thrypticocirrus rogickae* Hayward & Thorpe, 1988 | RS | 75º | - | - | 4.33 | - | 1 | IMC | Taylor et al. 2009 |
| *Valdemunitella* cf. *lata* (Kluge 1914) | CB | 62°09′ | 52°27′ | 70 | 4.77 | - | 1 | IMC | Krzeminska et al. 2016 |

**Table S3.** Q test and random effects model results for the meta-analysis.

| | | Q-Test Results | | Model Results |
|---|---|---|---|---|
| | df | Q | p | |
| *All Calcifiers Combined* | 24 | 48669.264 | < 0.0001 | Negative |
| *Aragonite* | 4 | 130.298 | < 0.0001 | Negative |
| *Calcite* | 6 | 39522.448 | < 0.0001 | Negative |
| *HMC* | 10 | 296.9688 | < 0.0001 | Negative |
| *LMC* | 1 | 0.6085 | 0.4354 | No effect |
